# The use of cross-validation has overestimated the value of genomic selection in plant breeding

**DOI:** 10.64898/2026.01.05.697784

**Authors:** Daniel E Runcie

## Abstract

Genomic Selection (GS) is widely considered to be a transformative approach for plant breeding, and has been a subject of well over a thousand papers since its proposal 25 years ago. The reduced costs of marker genotyping and genome sequencing, the proliferation of powerful statistical methods, and innovative breeding schemes that leverage GS have promised a revolution in the speed, efficiency, and precision of plant breeding. However, clear evidence of dramatically improved breeding outcomes using GS is difficult to find in the literature. I argue here that the most commonly presented evidence of GS success—high estimated accuracies of Genomic Prediction (GP) models as evaluated by crossvalidation—may be giving a highly misleading impression about the value of GS, at least in moderate-sized breeding programs. Estimating GP accuracy by cross-validation is only appropriate when GS is used to increase selection intensity, one of four key control parameters of the breeders equation and usually the least cost-effective way to increase genetic gain. If GS is instead used to increase the accuracy of selection among a fixed set of candidates or used to speed up breeding cycles, cross-validation-based estimates can be dramatically inaccurate, in ways that differ among breeding populations and traits. Instead, I show that analytical expressions and computational simulations are more informative about the likelihood of success of GS than cross-validation, and can be more effectively employed to evaluate GS program design.

## Introduction

The goal of plant breeding is to produce varieties with improved genetic values. Cycles of recurrent selection are the foundations plant breeding (Bernardo, 2010; Rutkoski, 2019). Each cycle involves generating a population of breeding candidates, evaluating them for genetic merit, selecting the best, and then recombining their alleles to generate a new population. Recurrent selection increases the average genetic value of the breeding population, which increases the probability of recovering a new best variety each generation. The increase in the average genetic value is called genetic gain and is usually optimized by selecting candidates based on their breeding value, which is twice the expected genetic value of their offspring when crossed randomly to other candidates in the population. The key factors that determine the rate of genetic gain, as described by the breeder’s equation (Comstock and Moll, 1963; Falconer et al., 1996), are the standard deviation of breeding values in the population (*σ*_*a*_), the accuracy of estimating these breeding values (*ρ*), the difference in mean estimated genetic values between the full and selected population (*i.e*, the selection intensity, *i*), and the speed of each cycle (1/*L*, with *L* the cycle length). The expected rate of genetic gain in a basic selection scheme is simply the product of these four terms: Δ_*h*_ = *i* · *ρ* · *σ* / *L*.

The rate-limiting step in traditional plant breeding schemes is typically the collection of phenotype data to estimate the breeding value of each candidate (Jannink et al., 2010; Cobb et al., 2013). Depending on the species and mating system, this may involve creating inbred lines from each candidate, growing them in multi-location, multi-year trials, and measuring traits like cooking time and nutritional quality that are time-consuming and expensive. The time and cost of phenotypic evaluation limits the number of candidates that can be evaluated (limiting intensity, *i*), the accuracy of their evaluation (*ρ*), and the speed of each cycle (1/*L*).

Genomic Selection (GS) can increase the rate of genetic gain in breeding in multiple ways by leveraging genome-wide marker data on breeding candidates. Genome-wide marker data can be used in a Genomic Prediction (GP) model to predict the breeding value of each candidate, which can then be used to increase accuracy, intensity, or speed (Meuwissen et al., 2001; Bernardo, 2010). GP models can be used to increase accuracy (*ρ*) by sharing phenotype information among relatives more effectively than with a pedigree by BLUP because genome-wide marker data can identify subtle relationships among founders and predict the effects of Mendelian segregation within families (Hayes et al., 2009; Habier et al., 2013). GP models can be used to increase selection intensity (*i*) by evaluating more candidates for the same total cost if genome-wide marker data is less expensive than phenotype data. Assuming a minimum number of candidates *k* must be selected to maintain genetic variation, evaluating a larger population *n* leads to higher intensity. GP models can also be used to increase the speed of each cycle if genome-wide marker data can be collected before phenotypic data is available, enabling earlier selections and crossing without waiting for inbred development, multi-environment trials, or wet-chemistry assays (Heffner et al., 2010). Beyond their use in GP models, genome-wide marker data can also be used to manage genetic variation within the breeding population (Jannink et al., 2010; Sonesson et al., 2012), and to optimize the shuffling of alleles through crossing decisions (Zhong and Jannink, 2007; Daetwyler et al., 2015). But I will focus on the application of Genomic Prediction models in Genomic Selection in this article.

Genomic Prediction models are statistical or Machine Learning tools that take in genetic marker data and output predictions of genetic values (Meuwissen et al., 2001; Jannink et al., 2010). Some GP models use additional features such as phenotype or phenomic data and environmental covariates to improve predictions. Forming predictions based on the genetic markers of a specific candidate requires a large number of parameters. Optimizing these parameters requires phenotypic data from a training, or reference (REF) population of individuals with genotype and phenotype data, and commonly some form of regularization to prevent over-fitting. By far the most commonly used GP architecture is the Genomic BLUP (GBLUP) model or the equivalent Ridge Regression BLUP (RRBLUP), although more complex architectures are sometimes used (VanRaden, 2008; Endelman, 2011; Jannink et al., 2010).

Regardless of its specific architecture, each GP model needs to be evaluated to confirm that it works sufficiently well to enable GS. For breeding, the most important evaluation metric of a GP model, as described by the breeder’s equation, is the Pearson’s Correlation Coefficient (PCC) between predicted genetic values and true breeding values among the current population of breeding candidates (*ρ*) (Falconer et al., 1996; Meuwissen et al., 2001; Jannink et al., 2010). This differs from metrics like the square Root of the Mean Squared Error (RMSE) of predictions, which evaluates the error of each individual candidate but is less relevant to the rate of genetic gain. Measuring the accuracy of a GP model is important for deciding among candidate GP architectures and for deciding whether Genomic Selection itself is likely to improve a plant breeding program over traditional phenotype-based selection.

There are two key challenges to measuring the accuracy of a GP model based on the above definition.

First, GP accuracy must be measured in a population of breeding candidates that is separate from the population used to train the GP model to avoid inflated estimates of accuracy. Since GP model accuracy is highest when the population used to train the model is closely related to the population of breeding candidates, GP model accuracy is usually measured by cross-validation, meaning that the breeding candidates are split into training and testing partitions, and accuracy is measured only in the testing partition (Legarra et al., 2008; Crossa et al., 2017). This has a few issues: i) The predictand, the true breeding values of each of the candidates, is not known. In animal breeding, GP models are often evaluated against estimated breeding values (EBVs) based on progeny tests with sufficient progeny that the EBV are measured with very little error (Legarra and Reverter, 2017, 2018), but this is very rare in plant breeding. Instead, GP predictions are usually evaluated against phenotypic data, so estimates must be corrected for the expected difference between phenotypic values and breeding values (Legarra et al., 2008; Ould Estaghvirou et al., 2013; Runcie and Cheng, 2019; Wang et al., 2025). ii) Given the uncertainty in the true breeding values, estimates of accuracy can be very noisy, a point I will illustrate further below. iii) Since phenotypic data are needed to train and test the GP model, and different training populations will result in different GP performance, breeders must commit to GS before knowing if it will work. To decide if GS will be beneficial, it would be better to be able to evaluate a GP model for different candidate breeding schemes prior to actually choosing a scheme and devoting years of time and resources to an approach that may not be successful (Dekkers et al., 2021). An alternative method to evaluate GP is by mathematical analysis. Several authors have proposed analytical expression for expected GP accuracy, although most assume unrealistic properties of training populations such as a lack of population structure (Daetwyler et al., 2008; Goddard, 2009; Hayes et al., 2009), but see (Dekkers et al., 2021), and rely on knowledge about genetic architecture such as heritability and the number of quantitative trait loci (QTL) and their linkage with genetic markers.

Second, the accuracy of a particular GP model will be different for every different population of breeding candidates. This means that there isn’t a single accuracy for a GP model, even if it is to be trained on a specific trait in a specific breeding program. That same model may have a high accuracy for one population of breeding candidates (*i.e*. target, or NEW population) but low accuracy for another. As described above, GP models can be used in three distinct ways to increase the rate of genetic gain: 1) increasing accuracy within a fixed population of candidates with phenotype data; 2) increasing intensity by evaluating a larger population of candidates; or 3) increasing speed by evaluating a new population of candidates that is too young to have been phenotyped. Each of these applications involves a different NEW population, thus the accuracy of a GP model will be different in every case. This means that reporting “the accuracy” of a GP model has little meaning. It is necessary to report the accuracy for a specific NEW population for a specific application of GS in a specific breeding program. As I illustrate below, this is not common practice in the current plant breeding literature.

Given these challenges with measuring GP accuracy, and the challenges and expense of implementing GS in moderate-sized plant breeding programs, critically evaluating the success of GS is important. The ultimate goals of GS are to increase the rate of genetic gain in time per cost, and to accelerate the release of improved varieties. There is now strong evidence that GS has improved cattle breeding programs (García-Ruiz et al., 2016; Doublet et al., 2019; Wiggans and Carrillo, 2022). However, evidence in plant breeding is limited, with existing studies showing at best moderate gains relative to phenotypic selection (Beyene et al., 2015; Rutkoski et al., 2015; Vivek et al., 2017; Zhang et al., 2017; Das et al., 2020; Bonnett et al., 2022; Butoto et al., 2022; Bandillo et al., 2023; Gesteiro et al., 2023). Measuring genetic gain is time-consuming and labor-intensive (Rutkoski, 2019), and unless controlled experiments with different breeding strategies are run in parallel, changes in gain caused by GS can be difficult to distinguish from changes caused by other factors such as different levels of investment, climate change, and changes in breeding objectives.

Therefore, I believe that the best empirical evidence that GS is actually transformative for plant breeding remains the estimates of GP accuracy from hundreds of academic papers. In the remainder of this paper, I aim to critically evaluate that conclusion. First, I show that the only potentially transformative application of Genomic Prediction models in plant breeding is to increase speed; that applications targeting increases in intensity and accuracy cannot greatly increase the rate of genetic gain. This observation is widely known, but perhaps not commonly quantified explicitly. Despite this result, I then show that the vast majority of the academic literature over the past 20 years has not studied GP accuracy in a way relevant to the goal of increasing speed. Instead, most papers report estimates of GP accuracy using cross-validation on diverse breeding populations in ways that can only provide insight into the use of GS to increase selection intensity, and rarely account for imprecise estimates of true breeding values. Since appropriate empirical data are rare, I show using simulations across a wide range of plausible genetic architectures that GP models trained on diverse breeding populations, with empirical estimates of accuracy similar to those in the literature, typically have very low accuracy when used to increase cycle speed. Using GS to increase speed requires applying GP models to individuals of future generations before those generations can be measured for phenotypes. I show that GP accuracy in full-sib populations derived from diverse populations tends to be dramatically lower than that estimated by cross-validation, and while GP accuracy can be higher in more diverse future populations, this does not translate into greater genetic gain. However, this discrepancy is not equal for all types of breeding populations, and I provide a novel technique to illustrate *in advance* whether a particular breeding population is likely to be useful as a starting point for GS. I conclude that the current evidence for a transformative success of GS in plant breeding in the academic literature is weak, and that the current literature standard that uses cross-validation to demonstrate GP models needs to be re-evaluated.

I wish to note here that similar criticisms of cross-validation and the different meanings and interpretations of Genomic Prediction accuracy have been made repeatedly in the plant breeding literature. In particular, Windhausen et al. (2012) and Riedelsheimer et al. (2013) both noted the typically (and theoretically expected) low within-family accuracy using diverse training populations within the context of recurrent Genomic Selection breeding schemes, and Werner et al. (2020) discussed the unreliability of cross-validation for predicting within-family accuracy in cases with population structure. Nevertheless, I believe that these papers did not go far enough in recommending the full re-evaluation of cross-validation as an evaluation metric for Genomic Selection, and so I aim to reiterate their points here.

## Methods

### Analysis of the sensitivity of genetic gain to different Genomic Selection approaches

The breeder’s equation can be written as Δ*z* = *i* · *ρ* · *σ*_*a*_ / *L* where Δ*z* is the rate of genetic gain, and the other parameters are defined in the Introduction. Genomic Prediction models can be used with Genomic Selection to increase the rate of gain by targeting three of these parameters: increasing intensity (*i*) by evaluating a larger population of breeding candidates; increasing accuracy (*ρ*) by better leveraging genetic relationships and Mendelian segregation; and reducing cycle lengths (*L*) by enabling selections prior to phenotyping. While real breeding programs differ from the idealized setting where the breeder’s equation applies (Rutkoski, 2019), a mathematical analysis of this equation can provide insight into the sensitivity of breeding outcomes to changes in breeding schemes. Here, I use analytical expressions for the expected increase in relative rates of genetic gain due to GS relative to phenotypic selection to study the sensitivity of gain to GS-based changes in these three parameters individually. Although it is unrealistic to assume that one parameter can be changed independently of the others while maintaining a fixed total cost, this sensitivity analysis provides insight into where GS could potentially have the most impact.

To improve intensity (*i*), Genomic Prediction models can be used to evaluate additional candidates from a broader breeding population that are genotyped but not phenotyped. If we assume that estimated genetic values of the candidates follow a normal distribution, the expected standardized selection intensity when *k* candidates are chosen from a pool of size *n* is (Bulmer, 1971):

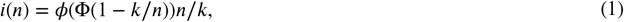

where *ϕ*(*x*) and Φ(*x*) are the pdf and cdf of the standard normal distribution, and I write *i*(*n*) to emphasize that *n* is the parameter that is controlled by increasing investment in GS. To explore the effect of investing in GS, I fixed *k* = 20 and assumed that the number of phenotyped candidates from the breeding population was *n*_*p*_ = 300, 600, or 2000, and then predicted the relative standardized selection intensity if an additional *n*_*g*_ = 1, …, 5000 candidates from the broader population were evaluated by the GP model: *i*(*n*_*g*_ + *n*_*p*_)/*i*(*n*_*p*_). The relative rate of genetic gain is expected to equal the relative selection intensity, assuming the other parameters of the breeder’s equation stay constant.

To improve accuracy (*ρ*), Genomic Prediction models use genomic data to refine estimates of breeding values made using the available phenotype data from each breeding candidate (Hayes et al., 2009; Habier et al., 2013). One strategy for approximating the accuracy of genomic predictions relative to phenotypic predictions or pedigree-based predictions is to use selection index theory to measure the impact of pedigree and genomic relationships. Dekkers et al. (2021) showed that Genomic Prediction accuracy for individuals with phenotype data can be approximated as:

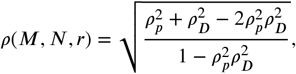

where 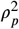 is the squared accuracy using only phenotypes and pedigree information and 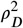 is the squared accuracy using the matrix **D** = **G** − **A**, with **A** the pedigree relationship matrix and **G** the estimated genomic relationship matrix among breeding candidates. 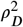 measures the information provided by genomic data that is not contained in the pedigree. I write *ρ*(*M, N, r*) as a function of *M*, the number of markers, *N*, the size of the phenotyped breeding population, and *r*, the number of phenotypic replicates per candidate, because for any given breeding population, the effectiveness of GS will depend on the investment in each of these parameters.

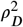 itself can be predicted as:

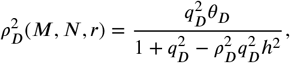

with 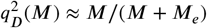 (Goddard et al., 2011), the proportion of genetic variance captured by markers, assuming markers and the true quantitative trait loci (QTL) have the same distribution and that *M*_*e*_ represents the effective number of independently segregating chromosomal segments as defined by (Visscher et al., 2006), *h*^2^ the narrow-sense heritability of the trait, and 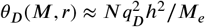.

To explore the effect of investing in GS, I defined four contrasting breeding populations differing in *h*^2^, *ρ*_*p*_, and *M*_*e*_: Two populations were diverse (*M*_*e*_ = 3700, similar to the estimate from a chicken population by Dekkers et al. (2021)), and two populations had a narrow genetic basis like a biparental population (*M*_*e*_ = 59, the average estimate from maize biparental populations by Lian et al. (2014)); Two populations had high heritability (*h*^2^ = 0.4) and two had low heritability (*h*^2^ = 0.1); The two diverse populations had additional information from related individuals through a pedigree 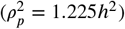, while the two narrow populations had no pedigree information; While not necessarily representative of any real breeding populations, these parameter choices were designed to span realistic ranges for plant breeding populations while illustrating the impact of each parameter on the relative rates of genetic gain due to GS.

Using each of these breeding populations, I measured the relative rates of genetic gain (relative to phenotypic selection) as *M* increased from 100 to 100,000, as *N* increased from 300 to 30,000, or as *r* increased from 1 plot per candidate to 8 plots per candidate (with *h*^2^ calibrated at the above values using 2 plots per candidate). For studying the impact of *r*, I assumed that the additional information from relatives through the pedigree 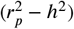 was constant as a function of *r*, and all genetic variation was additive, so plot means converged to breeding values with increasing *r*.

To improve speed (1/*L*), one or more selection cycles can be based entirely on Genomic Predictions. For example, if *Y*_*p*_ years are needed for a phenotypic selection cycle (*e.g*., to generate inbred lines, make top-crosses, and run a sufficiently large multi-environment trial), and then *Y*_*g*_ years are used to run *N*_*g*_ Genomic Selection cycles, the relative rate of gain is *Y*_*p*_(1 + *N*_*g*_ )/(*Y*_*p*_ + *Y*_*g*_ ). I set *Y*_*p*_ = 5, *Y*_*g*_ = 0 − 2, and *N*_*g*_ = 1, …, 8, assuming that four rapid cycle generations could be run per year, either for inbreeding and seed bulking, or for rapid cycle GS, three years would be needed for a multi-environment trial, and that if 1+ GS cycles were used, the phenotypic cycles would be delayed by 1-2 years. Note that this is a simplified scheme where a new GP model is trained each phenotypic selection cycle. More complex schemes with continually up-dating training data could potentially increase speed even more.

It is also possible to improve speed by allowing selection on the current breeding population earlier; for example if phenotypic selection requires 5 years for inbred development, multi-environmental trials, and wet-chemistry assays, but GS could be applied after 4 years using only one year of trials. I discuss the potential benefit of this approach below.

### Literature survey

Using PubMed, I searched for journal articles with the terms “Genomic Selection OR Genomic Prediction” AND “plant OR Crop” in the title and/or abstract with publication year through 2024. Of the 1,022 papers published since 2009 returned by this search, I randomly screened for ones that trained at least one Genomic Prediction model on real phenotype data from plants, drawing new papers from the search results until I had a collection of 20 relevant papers per 5-year period (except I failed to find enough in the 2009-2012 period and ended up with only 5 in this period). For each paper, I scored 1) whether or not Genomic Prediction accuracy was measured using cross-validation, using analytical predictions of accuracy, or not evaluated, and 2) if the accuracy was measured in partitions of the same population used to train the model or in a different population, and if a different population, whether that population was derived from the population used to train the model, and thus could be considered a future population that could not have phenotype data available at the time that the Genomic Prediction model was trained. I also assessed whether the training and prediction populations were diverse and if they were likely to exhibit population structure.

For each paper meeting the above criteria, I extracted a representative estimate of Genomic Prediction accuracy. Many papers report multiple estimates of accuracy—for different traits, different models, different environments, etc. When a paper evaluated multiple models, I extracted accuracy measures for GBLUP or snpBLUP or RRBLUP if it was included, otherwise the best model. When a paper evaluated models for multiple traits, I extracted accuracy measurements for yield if that was included, otherwise the first trait reported if presented sequentially, or the average of all traits reported together. I excluded CV2 and CV0-style genomic predictions, focusing only on accuracy for candidates without any phenotype data, as these have more clarity in their value for Genomic Selection, for reasons I discuss below. When the first report of accuracy meeting the above criteria included a set of conditions (environments, traits) all reported together, I extracted the average if available, or the (max + min)/2 if given as a range. For each accuracy estimate, I recorded the size of the training population, whether the training population was diverse or a narrow family (fewer than 10 founders), the average accuracy estimate, the species, and whether the accuracy was estimated as “predictive ability” (*i.e*. Pearson’s Correlation Coefficient with phenotypic value), or “predictive accuracy” (*i.e*. Pearson’s Correlation Coefficient with the breeding value), which is usually estimated by dividing a predictive ability estimate by the square root of the trait heritability (Legarra et al., 2008; Ould Estaghvirou et al., 2013).

### Simulation Study

I used simulations to explore the reliability of cross-validation as an estimate of the accuracy of a Genomic Prediction model in a Recurrent Genomic Selection breeding scheme. Simulations were designed to span both factors that a breeder can control or observe (size and diversity of the REF population, and the estimated narrow-sense heritability of the trait), and factors that they cannot (genetic architecture).

To ground simulations in a relevant plant breeding context, I used maize genetic marker data from the GenomesToFields initiative (Lima et al., 2023).

### Selection of six REF populations

I downloaded genetic marker data for the GenomesToFields maize hybrids from Lima et al. (2023). Since phased genotype data are needed to simulate offspring genotypes, I phased the genotypes of each hybrid using the provided pedigree. Specifically, I collected all hybrids with the same male parent (Hybrid_Parent2), then inferred the inbred genotypes of the female parents using the following rules:

- If all hybrids had genotype 0/0 or 0/1 (meaning the male parent had genotype 0/0), the female parent must be 0/0 for a hybrid with genotype 0/0 and must be 1/1 for a hybrid with genotype 0/1.
- If all hybrids had genotype 0/1 or 1/1 (meaning the male parent had genotype 1/1), the female parent must be 0/0 for a hybrid with genotype 0/1 and must be 1/1 for a hybrid with genotype 1/1.

Markers that did not fit this pattern were assumed to be genotyping errors and were assigned NA, and then inputed as 0/0 or 1/1 randomly based on the frequency of the 1 allele in the whole population. For inbreds contributing to more than a single hybrid, I selected the genotype constructed as above with the fewest missing values.

Missing hybrid genotypes were imputed as 0/1/2 based on rounding twice the population allele frequency to the nearest integer. After filtering, the final datasets included hybrid genotype data for 4,928 hybrids at 427,478 markers and inbred genotype data for 2,190 inbreds, including all parents of the 4,928 hybrids.

From the 4,928 hybrids, I selected six subsets to represent a range of reference (REF) populations:

1. “Biparental_212”: The 212 hybrids of the PHN11xPHW65 biparental population crossed to the LH195 tester
2. “MAGIC_387”:The 387 hybrids from the WI-MAGIC population crossed to the PHZ51 tester.
3. “NAM_566”:The 566 hybrids of the PHN11xPHW65, MO44xPHW65, and PHW65xMOG biparental populations, all crossed to the LH195 tester. This forms a small nested association mapping population with the PHW65 genotype as the recurrent parent.
4. “Diversity_212”: 212 randomly selected hybrids excluding the above hybrids as well as hybrids using the same female parents.
5. “Diversity_387”: 387 randomly selected hybrids excluding the above hybrids as well as hybrids using the same female parents.
6. “Diversity_566”: 566 randomly selected hybrids excluding the above hybrids as well as hybrids using the same female parents.

### Simulating genetic architecture

I designed five contrasting classes of genetic architecture varying to different degrees from the architecture assumed by the GBLUP and snpBLUP models.

1. “simple_additive”: I sampled 5,000 SNPs randomly from all available SNPs, and drew additive effect sizes for each from a standard normal distribution.
2. “complex_additive”: Same as “simple_additive”, but using 100,000 SNPs randomly from all available SNPs.
3. “rare_additive”: Same as “simple_additive”, but using 27,872 markers with MAF < 0.01 across the whole hybrid population (excluding the NAM and WI-MAGIC populations). Note that since the SNPs in the GenomeToFields dataset were imputed from a Practical Haplotype Graph with 86 founders (Washburn 2024), rare variants are underrepresented in these QTL sets, so the other two additive architectures are biased towards common variants.
4. “simple_dominance”: Using the same 5,000 SNPs as the “simple_additive” architecture, I created an architecture with both additive and dominance variance following the approach of Wellmann and Bennewitz (2011), using the parameters from Scenario 3 (Caballero and Keightley, 1994). Specifically, I sampled dominance coefficients: *δ*_*j*_ ∼ N(0.2, 0.3), additive effects: *a*_*j*_ ∼ N(0, *exp*(3*δ*_*i*_)), and then set the dominance effects to: *d*_*j*_ = *δ*_*j*_ |*a*_*j*_| . I then re-assigned *a*_*j*_ = |*a*_*j*_| sign(1 − 2*q*_*j*_ ), for *q*_*j*_ the frequency of the alternate allele in the full population. This ensures that the magnitude of dominance and additive effects is positively correlated, and that deleterious alleles tend to be rare.
5. “rare_dominance”: Same as “simple_dominance”, but using the 27,872 rare SNPs.

For each set of additive and dominance SNP effects, I calculated the genotypic effect for each individual as *g* = ∑_*j*_ *a*_*j*_ *x*_*i*_ +*d*_*j*_ *I*(*x*_*j*_ = 1), for *x*_*j*_ ∈ 0, 1, 2 the number of copies of the alternative allele in individual *i*. I then rescaled the additive and dominance effects so that var(*g*) = 1. Finally, I calculated the allelic substitution effects of each locus as *α*_*j*_ = *a*_*j*_ + (1 − 2*q*_*j*_ )*d*_*j*_, and the breeding value of each individual as ∑_*j*_ *α*_*j*_ *x*_*j*_

To complement the genetic architecture, I sampled an additional set of 20,000 SNPs as genetic markers.

### Simulating phenotype data to calibrate cross-validation accuracy estimates

For each simulation, using each architecture, I simulated phenotype data for the REF population for a single trait by adding random environmental noise to the genetic value of each hybrid so that the cross-validation-based estimate of GBLUP predictive ability would equal a predefined value: *r*_0_ = 0.3, 0.5, or 0.7.

First, I drew a vector (**e**) of random independent environmental noise for each individual in the REF population, and added this to the genotypic vector to form candidate phenotypes: **y** = **g** + **e**.

I then divided the REF population into 5 non-overlapping folds and for each fold, I fit the following GBLUP model using the *mixed.solve* function of the `rrBLUP` *R* package (Endelman, 2011) to the four remaining training partitions:

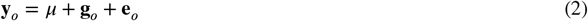

where 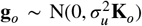 are the genetic values of the training (“old”) data, 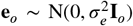, and 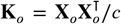 with **X**_*o*_ the centered genotype matrix from the 20,000 SNP markers in the training data. Predictions for the test data were formed as 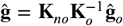, where 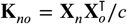, with **X** the genotype matrix from the same SNP markers in the testing (NEW) data, centered by the same values as **X**_*o*_. Predictive abilities 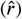 were recorded as the average Pearson’s Correlation Coefficient PCC between predicted genetic values 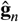 and observed phenotype data **y**_*n*_ over the 5 partitions.

I then used the ‘optimize’ R function to optimize a scale parameter *c* to multiply the error vector to increase or decrease the magnitude of environmental error until the difference between the estimated predictive ability and the target was 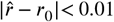: **y** = **g** + *c***e**.

### Measuring GBLUP accuracy

After calibrating the phenotype data for the REF population as above, I fit the GBLUP model (2) to the full REF population. I then measured the actual prediction accuracy of that model for three relevant NEW populations:

1. The REF population itself. This is relevant to using Genomic Selection to improve selection accuracy.
2. A set of 2533 additional hybrids from the GenomeToFields project. This is relevant to using Genomic Selection to increase selection intensity.
3. A F2 population derived from the non-tester inbred parents of two hybrids selected from the REF population based on their phenotypes. This is relevant to using Genomic Selection to increase speed. I selected two hybrids randomly from those in the top 10% of phenotypic values and then simulated 500 F2 offspring from their cross, using AlphaSimR (Faux et al., 2016; Gaynor et al., 2021) and a genetic map from Ogut et al. (2015).

In each each case, I measured prediction accuracy as 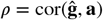, with **a** = ∑_*j*_ *α*_*j*_ **x**_*j*_ the breeding value for each individual.

### Predictions of Genomic Prediction accuracy

As an alternative to cross-validation, I used the following formula to predict GBLUP accuracy for each NEW population:

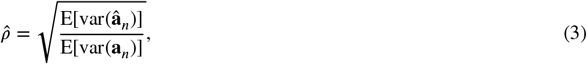

where **a**_*n*_ = **X**_*n*_***α*** are the true additive genetic values of the NEW candidates, **X**_*n*_ is the matrix of QTL genotypes coded as {0, 1, 2} − 2*q*, with *q*_j_ the minor allele frequency of the *j*th QTL in the REF population, and 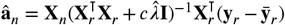 would be the GBLUP predictions if the markers were identical to the QTL themselves. Note that in fact, Markers and QTL were non-overlapping sets of variants as described above. var(**a**_*n*_) and var(*â*_*n*_) are the sample variances of the true and predicted additive genetic values, and the expectations are taken with respect to the unknown QTL effects (**b**) and as yet un-measured phenotypes 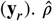 can be calculated for hypothetical NEW populations before phenotype data are collected on the REF population as long as an estimate of 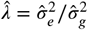, the ratio of estimated residual and additive genetic variances is available. I used the estimate of this value from the GBLUP model fit to the phenotypes of the REF data. However, in principle, values from prior experience with the same traits in similar populations could be used so that this value could be estimated before the REF population was phenotyped.

As I show in the Supplemental Materials, once **y** is observed on the REF population, an alternative expression can be used which is expected to be more precise and less sensitive to departures from model assumptions. These two expressions are derived in the Appendix and are based on the assumptions that the markers are the QTL, that the genetic and residual variances are known, and that the the NEW candidates are derived from the REF population without selection. I test the impact of violations of these assumptions in the simulation studies. In the Appendix, I also derive a simplified form of Equation 3 using the singular value decomposition of the REF population genotype matrix 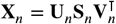, which provides an intuitive explanation for why Recurrent Genomic Selection will work better for some REF populations than others.

### Data availability

Genotype data underlying the simulations was downloaded from https://datacommons.cyverse.org/browse/iplant/home/shared/commons_repo/curated/GenomesToFields_GenotypeByEnvironment_PredictionCompetition_2023. Scripts to reproduce all analyses are available on Github: https://github.com/deruncie/Predicting_GP_accuracy.

## Results and Discussion

### Genomic Prediction models can be dramatically more effective if used to increase speed rather than increase selection intensity or accuracy

Genomic Selection strategies to increase intensity (*i*), accuracy (*ρ*), or reduce selection time (Speed, 1/*L*) require applying Genomic Prediction models to distinct populations of candidates in a breeding program (Figure 1A). To explore the potential benefits of each application of Genomic Selection, I applied analytical formulas to predict the impact of investing in genotyping and Genomic Prediction.

**FIGURE 1.**
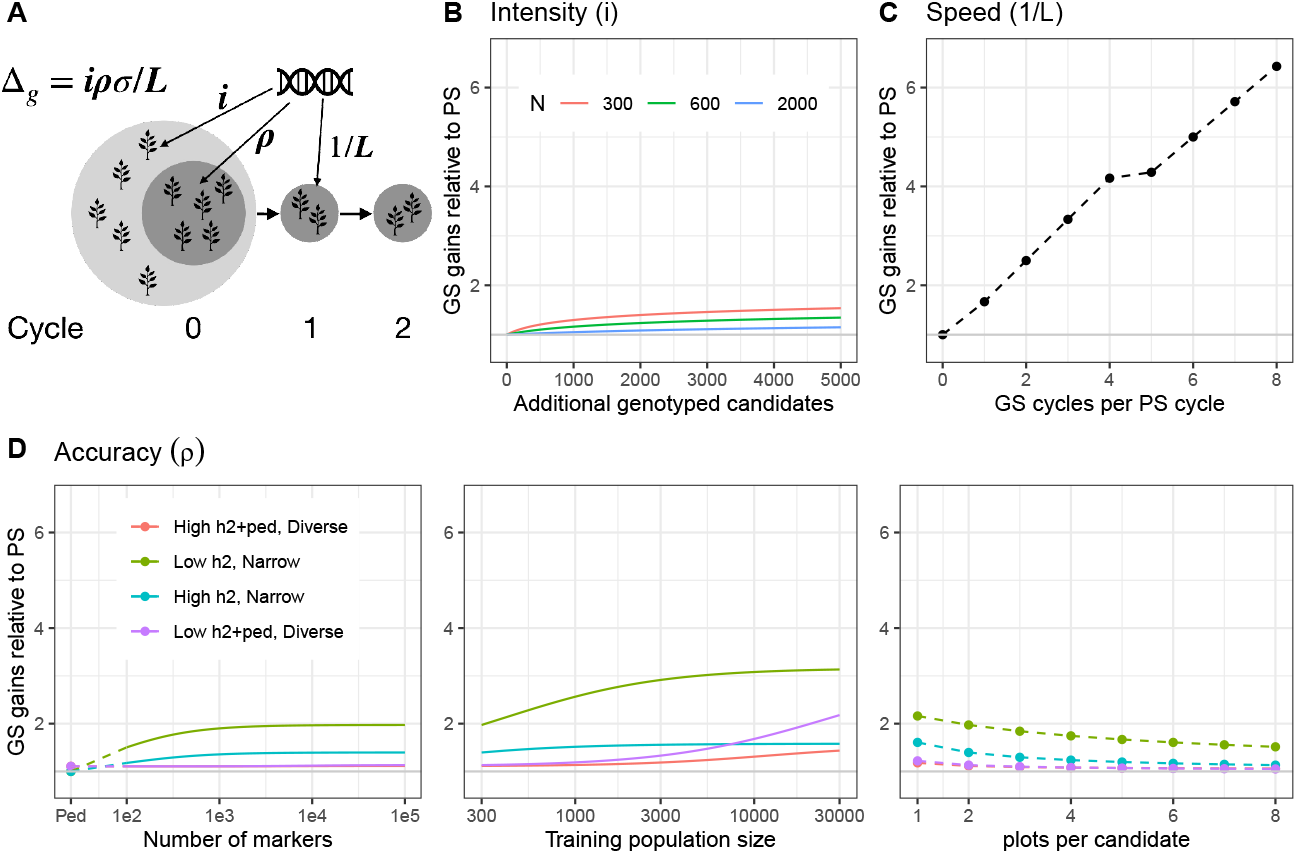
An analysis of the breeder’s equation illustrates why the most impactful use of Genomic Prediction is to enable faster recurrent selection cycles. **A**. Illustration of a recurrent breeding scheme and opportunities to use a Genomic Prediction (GP) model for selections by targeting distinct populations of selection candidates. Each of the subsequent panels evaluates the expected ratio of genetic gains based on the breeder’s equation if Genomic Selection was used to improve that specific parameter, relative to a phenotypic selection scheme. **B**. Rates of genetic gain when using GP to evaluate additional candidates of the Cycle 0 generation without phenotype data, relative to phenotypic selection for three sizes of populations of phenotyped candidates as a function of the number of additional candidates evaluated by GP. **C**. Rates of genetic gain when using GP to evaluate candidates of future cycles, eliminating the need to wait for phenotype data, *i.e. Recurrent Genomic Selection*, relative to phenotypic selection. Values assume PS cycles require 5 years and GS cycles can occur 4x per year. **D**. Rates of genetic gain when using GP to improve the accuracy of the reference population of candidates in Cycle 0, *i.e*., those with phenotype data, relative to phenotypic selections of these candidates. Three ways to increase the intensity of GP are evaluated: i) increasing the number of markers genotyped, ii) increasing the reference population size, and iii) improving the quality of the phenotype data by higher replication per candidate. In each panel, four types of reference populations are used, varying in genetic diversity (Diverse vs Narrow), heritability (*h*^2^), and whether a pedigree is available and informative (+ped). Exact parameters are given in the Methods. A zoomed-in version of the figure is available as Supplemental Figure S1.

The gain from using Genomic Prediction to increase selection intensity in a particular breeding cycle depends on the size of the population of candidates that could be selected by phenotype alone and the number of additional candidates that can be evaluated by Genomic Prediction. Assuming selection accuracy is the same for these two groups, adding up to 5000 additional candidates would result in gains due to GS of up to ≈ 50% if starting from 300 phenotyped candidates, but only up to ≈ 15% if starting from 2000 phenotyped candidates (Figure 1B, see Supplemental Figure S1B for zoomed in version). If only 500 additional candidates are genotypyed, maximum gains are ≈ 21% and ≈ 3%, respectively.

The gain from using Genomic Prediction to increase the selection accuracy of a fixed pool of candidates that has phenotype data depends on several factors, including properties of the candidates themselves (*N, h*^2^, genetic diversity, pedigree relationships), as well as the number of markers used for Genomic Prediction. Genomic Prediction can increase accuracy up to ≈ 3*x*, but only for large populations with a very narrow genetic basis (*e.g*., biparental maize populations, *M*_*e*_ = 59, based on maize biparentals (Lian et al., 2014)) and for traits with low heritability (Figure 1D, Supplemental Figure S1D). For a moderately diverse population (*M*_*e*_ = 3700, based on estimates from a chicken population (Dekkers et al., 2021)), increases in accuracy reached ≈ 2*x*, but only for a population size of 30,000 individuals. For a population size of 300, more typical of many plant breeding programs, increases in gain were < 15%, but almost all of that was due to phenotypic information from relatives through the pedigree (which is also captured by Genomic Prediction (Habier et al., 2013)). Additional gains from Genomic Prediction beyond the pedigree relationships were only ≈ 2% and ≈ 1% for low and high *h*^2^, respectively. For fixed population sizes, gains from increasing the number of markers used in Genomic Prediction saturated earlier for the narrow populations than the diverse populations, and increasing the phenotyping investment per candidate reduced the relative gains due to Genomic Prediction, even though this would make the Genomic Prediction models more accurate.

Compared to these two results, the gains from using Genomic Prediction to increase speed by reducing cycle times, commonly called “Recurrent Genomic Selection” were much more dramatic (Figure 1C, Supplemental Figure S1C), reaching > 6*x*. I assumed that for a Genomic Selection-based scheme, one cycle of phenotypic selection (five years) would be required to train a Genomic Prediction model, and then that model could be used for 1-8 cycles with selection purely on genomic-predicted breeding values which each could advance rapidly using speed breeding (4/year) since no phenotype data would be needed. Compared to potential gains from targeting intensity or accuracy in a diverse population with moderate heritability (*N* = 300, *h*^2^ = 0.4), the potential gains from speed due to Genomic Selection as modeled here are > 800*x*.

It is also possible to improve speed by allowing selection on the current breeding population earlier; for example if phenotypic selection requires 5 years for inbred development, multi-environmental trials, and wet-chemistry assays, but GS could be applied after 4 years using only one year of trials, speed gains would equal 1.25*x*. However, applying GS earlier means that the candidates would have lower-quality phenotype data (Supplemental Figure S1A). Genomic Predictions would be expected to compensate for this loss of phenotypic accuracy following Figure 1D and Supplemental Figure S1D, so the total impact on genetic gain could be predicted as the product of these three effects. Using Supplemental Figure S1A, we see that if *h*^2^ = 0.4, a 50% decrease in phenotyping intensity would approximately cancel the gains from increasing speed from 5 years to 4 per cycle, so GS gains would come entirely from accuracy, following Supplemental Figure S1D. If *h*^2^ was higher, loss of phenotypic accuracy from reduced phenotyping would have less of a cost, so GS gains would be positive, but in contrast, if *h*^2^ = 0.3, less of the gains from accuracy would translate into total genetic gain. Nevertheless, this analysis suggests that enabling selections earlier within a phenotypic selection cycle could have at best a moderate effect on the rate of genetic gain.

However, it is critical to note that these analyses are *upper-bounds* for the potential increases in genetic gain from Genomic Selection and assume that implementing Genomic Selection to target one parameter can be done without impacting any of the other parameters of the breeder’s equation. This is clearly not realistic for several reasons: First, creating and genotyping additional candidates to increase intensity or run additional cycles imposes a significant cost on breeding programs. With a fixed budget, this will likely reduce the resources for phenotyping, likely reducing *h*^2^ and thus Genomic Prediction model accuracy. Second, unlike phenotyped candidates, candidates evaluated only by genomic-predicted breeding values will likely be evaluated less accurately. This is especially true for candidates in future cycles, because the accuracy of Genomic Prediction declines as the relationship between reference (REF) and target (NEW) populations diverge (Dekkers et al., 2021)). Thus the gains in Figure 1 (particularly B and C) are likely optimistic. The low predicted gains from Genomic Prediction in small, diverse populations (second panel of Figure 1D) are particularly concerning, as this indicates that such Genomic Prediction models are likely to have very low accuracy.

This uncertainty means that empirical measures of Genomic Prediction accuracy *as implemented in each of the three cases in Figure 1A* are critical. Potential increases in gain of only 2% − 21% by targeting accuracy, intensity, or within-cycle speed in diverse, moderate-sized breeding programs may be insufficient to justify the costs of Genomic Selection, notwithstanding that the actual numbers are likely to be lower. Increases in gain of up to 6*x* by targeting speed in Recurrent Genomic Selection would certainly be transformative to any breeding program. But these gains would only be possible if Genomic Prediction models remain accurate for many future generations. Since targeting speed is by far the most potentially transformative application of Genomic Prediction, I conducted a literature survey to assess the evidence that Genomic Prediction accuracy was sufficient in to enable greatly increased speed.

### Few studies of Genomic Prediction in plants have measured Genomic Prediction accuracy in a relevant Recurrent Genomic Selection context

Of the 65 randomly sampled papers reporting empirical evaluations of Genomic Prediction models in plant populations from 2009-2024, all evaluated Genomic Prediction performance using cross validation, but only 5 measured Genomic Prediction accuracy in a population without structure (Figure 2). Seven also calculated (6) or directly estimated (1) genetic gain across cycles. Only 21/65 designed cross-validation training and testing populations in a manner respecting the structure of existing genetic groupings, and of these, only 7 explicitly used NEW populations that could be considered “future” populations relative to the training population. Thus, the vast majority of studies report estimates that are directly relevant to using Genomic Prediction to increase intensity, but few report estimates relevant to using Genomic Prediction to increase speed.

**FIGURE 2.**
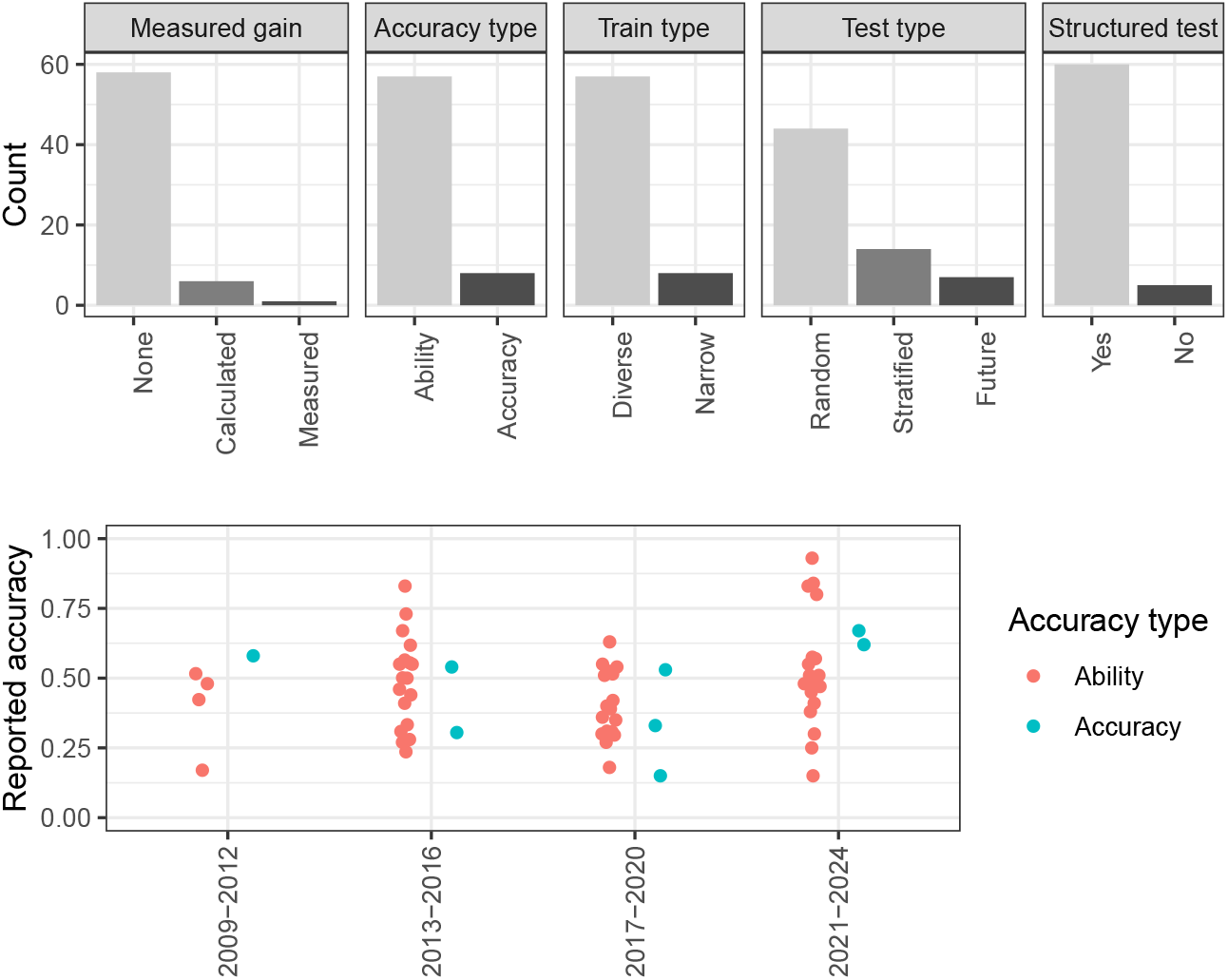
Published reports of Genomic Prediction accuracy in plant populations are typically high. However, papers usually report cross-validation estimates of *Predictive Ability* (Legarra et al., 2008; Ould Estaghvirou et al., 2013), and rarely use test populations without population structure. Recurrent Genomic Selection requires moderate Genomic Prediction accuracy within families and thus it is unclear if these published estimates are useful for predicting the success of Recurrent Genomic Selection.

Furthermore, 57/65 used diverse REF populations with sizes ranging from 70-2580 (Median = 304, Mean = 480), and only 8/65 estimated GP accuracy (i.e., Pearson’s Correlation Coefficient between predicted and true genetic values) instead of *ability* (i.e., the correlation with observed phenotypes), even though only the former is directly relevant to the response to selection in the breeder’s equation (Legarra et al., 2008; Ould Estaghvirou et al., 2013). No papers reported analytical predictions of Genomic Prediction accuracy for any population.

From each paper, I extracted a representative estimate of Genomic Prediction accuracy. Accuracy estimates varied widely across studies, but without a clear patterns across the year of publication, species, trait, or Genomic Prediction model used. The only pattern with a significant (p<0.05) association with reported accuracy was whether the testing population was genetically stratified from the training population. Estimates of accuracy were generally quite high. The overall average was close to *ρ* = 0.5, with 75% > 0.33 and 50% > 0.48.

If we take these estimates as representative of typical accuracies of Genomic Prediction models in plant breeding contexts, breeders should be able to run very short breeding cycles without waiting for phenotype data to make selections, and thus make tremendous gains in target traits. However, since the vast majority of these accuracy estimates were made in populations with population structure, and populations that do not accurately represent the genetic relationships between future generations and the populations used to train a Genomic Prediction model, it is unclear whether these accuracy estimates are meaningful for evaluating the potential of Recurrent Genomic Selection.

### Cross-validation estimates or Genomic Prediction accuracy are unreliable for evaluating Recurrent Genomic Selection

To study this question, I designed a simulation study to assess the potential discrepancy between cross-validation-based estimates of Genomic Prediction accuracy within random partitions of a reference population, and potential candidates in a future generation. I ran a total of 9,000 simulations, using genetic marker data from maize extracted from the GenomesToFields dataset, and simulated phenotype data for a single target trait. These simulations spanned six reference populations with different structures and five classes of genetic architecture, and were each calibrated so that the estimate of predictive ability (as this is the statistic most commonly reported, Figure 2A) equaled 0.3, 0.5, or 0.7.

Starting with a setting that approximately matches experimental designs and results from the literature: a diverse REF population of *N* = 387 hybrids and a trait with Genomic Prediction predictive ability using GBLUP estimated to be *r* = 0.5, I measured the actual accuracy of the GBLUP model when used to predict breeding values of i) the parents of the 387 hybrids used to train the model, *i.e*., reflecting the case where Genomic Prediction is used to increase accuracy; ii) a broader population of 2533 hybrids sampled from the whole GenomesToFields population, but excluding the large biparental and NAM families *i.e*., reflecting the case where Genomic Prediction is used to increase intensity; and iii) among 500 F2 plants created by simulating the offspring of a cross between two randomly selected parents from the top 10% of the predicted breeding values of the REF population, *i.e*., reflecting the case where Genomic Prediction is used to increase speed by making selections in the second generation. For the case of using Genomic Prediction to increase speed, I only considered prediction accuracy among the full-sib offspring of a single biparental cross. As I show in the Supplemental Materials, the precision of genomic prediction may be higher in “future” NEW populations that are derived from multiple full sib families, but this accuracy does not translate into genetic gain, a point also demonstrated by (Müller et al., 2017) and (Legarra and Reverter, 2017). Only within-family accuracy, or more generally, accuracy measured in NEW populations without structure, directly translates into improvements in total genetic gain after multiple cycles of recurrent Genomic Selection. I ran 100 simulations using each of five classes of genetic architectures, since the actual genetic architecture is never known in advance, and is very difficult to estimate empirically (de Los Campos et al., 2015).

In nearly every simulation, if targeting intensity or accuracy, the GBLUP model had much higher accuracy than estimated by cross-validation (Figure 3A). But if the model was used to increase speed, the actual accuracy of the GBLUP model was usually much lower. For additive genetic architectures, the true accuracy averaged ≈ 0.3 if driven by common variants, but only ≈ 0.15 if driven by rare variants. In ≈ 25% of cases, the true accuracy was less than 0.1. More importantly, the range of actual prediction accuracies varied widely, indicating that the cross-validation estimate of *r* = 0.5 was largely uninformative as to whether accuracy would be high or close to zero. For architectures with dominance variance, true accuracies varied even more widely.

**FIGURE 3.**
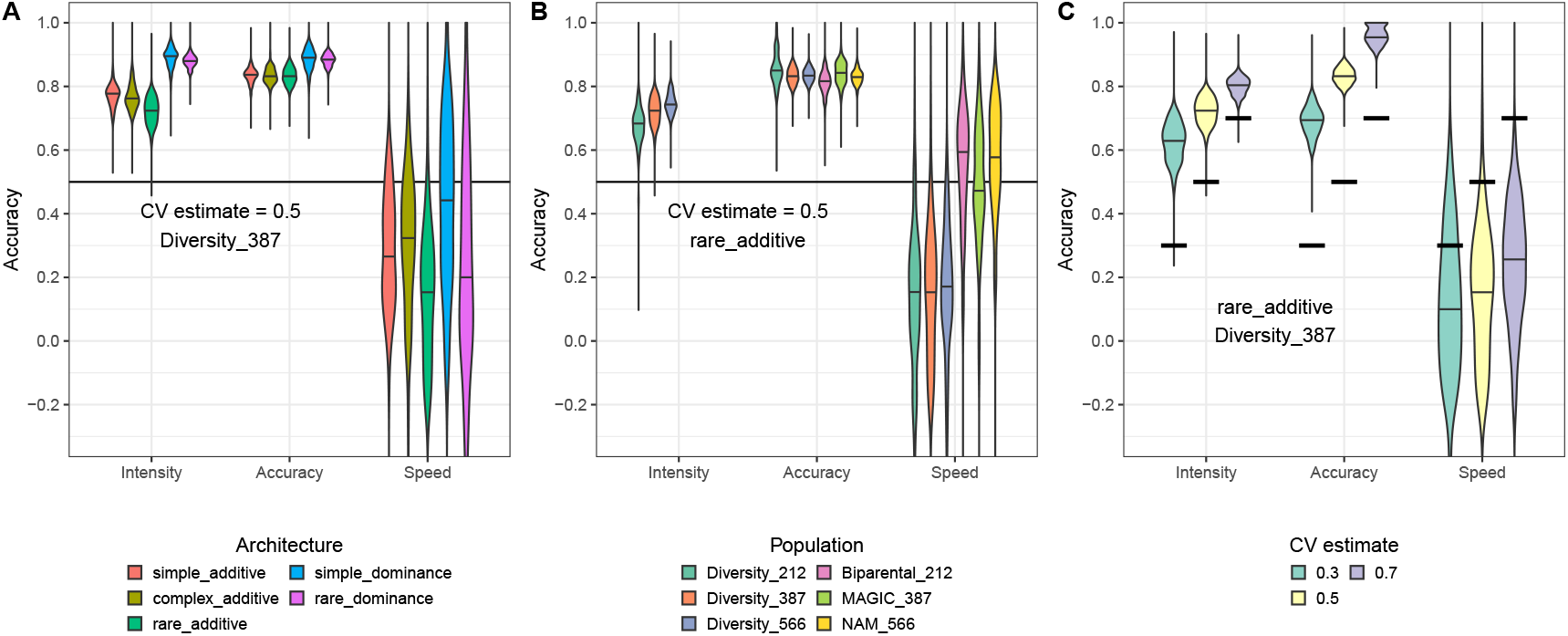
Cross-validation estimates of accuracy are unreliable for evaluating Genomic Prediction accuracy. Each plot shows the density of true accuracy estimates for NEW populations relevant to increasing intensity, accuracy, or speed. Violin plots show densities of 100 simulations per setting. Heavy black bars show the cross-validation predictive ability estimates. **A**. Simulations for the Diverse_387 population when cross-validation estimates predictive ability as *r* = 0.5 for five different genetic architectures. **B**. Simulations for six different REF populations when cross-validation estimates predictive ability as *r* = 0.5 using the `rare_additive`genetic architecture. Note that for the three narrow populations, no additional relevant hybrids exist, so it is not useful to use GBLUP to increase intensity. **C**. Simulations for the Diverse_387 population with a `rare_additive`genetic architecture, for different estimates of predictive ability by cross-validation.

I then repeated these simulations for five other REF populations: two additional diverse REF populations of size *N* = 212 and *N* = 566, and three narrow populations: a biparental population of size *N* = 212, a MAGIC population of size *N* = 387, and a NAM population of size = *N* = 566. Figure 3B shows results for simulations of a trait with a cross-validation-based estimate of predictive ability set to *r* = 0.5 and an additive genetic architecture driven by rare variants. Again, cross-validation significantly underestimated the true accuracy of GBLUP models if they were used to increase accuracy or intensity. However, results for increasing speed were mixed. For the three narrow populations, the cross-validation-based estimates were approximately unbiased.

But for the three diverse populations, the cross-validation-based estimates again significantly over-estimated the true accuracy. As in Figure 3A, though, more important than the bias is the lack of precision: observing *r* = 0.5 using cross-validation provides very little information about the actual GBLUP accuracy if used to increase speed.

This result is perhaps most clearly illustrated by Figure 3C, which compares the actual accuracies between simulations with cross-validation estimates of *r* = 0.3, 0.5, and 0.7 for the original diverse population of size *N* = 387 with a `rare_additive`genetic architecture. If cross-validation provided useful information about GBLUP accuracy, actual accuracies should be correlated with the cross-validation estimates. This is the case when GBLUP is used to increase accuracy or intensity: while cross-validation estimates are biased, the distributions of true accuracies are clearly different between traits with different estimates of predictive ability. But if GBLUP is used to increase speed, the distributions of true accuracies are largely indistinguishable. This demonstrates again that cross-validation provides very little information about the actual GBLUP accuracy if used to increase speed.

Results for the other four architectures, across the six populations, and three cross-validation-based estimates, are provided in Supplemental Figure S3. Additionally, I show that correcting cross-validation estimates by dividing by the square-root of estimated heritability to estimate prediction accuracy instead of ability (Legarra et al., 2008) results in less bias, particularly for intensity, but similarly dispersed estimates that are largely uninformative for speed.

To conclude, cross-validation estimates of the “predictive ability” of a GBLUP model are never accurate estimates of the actual accuracy of that GBLUP model when used for Genomic Selection. If the GBLUP model is to be used to increase selection intensity or accuracy, the cross-validation estimates are informative, but severely biased downwards in a way that is somewhat sensitive to the (unknown) genetic architecture and (known) characteristics of the REF population. However, if the GBLUP model is to be used to increase cycle speed, as in Recurrent Genomic Selection, the cross-validation estimates are not only biased in ways that vary significantly according to genetic architecture and REF population structure, but are also so imprecise that they provide very little information.

### Analytical calculations are more useful than cross validation for predicting the usefulness of Recurrent Genomic Selection

The simulations above demonstrate that cross-validation cannot reliably evaluate Genomic Prediction models, especially when they are used to increase speed using Genomic Selection. But even if cross-validation was accurate, and could indicate that Genomic Selection will work in some breeding populations, cross-validation would have limited utility because it requires phenotype data on the REF population. This means that a breeder must create a specific REF population and subject it to high-quality field trials before knowing whether or not it will work as as a training population for Genomic Selection.

It would be more useful if a candidate REF population could be evaluated for use in Genomic Selection from its genotypes alone. As I show in the Supplemental Materials, the expected correlation between predicted and true genetic values can be calculated for any pair of REF and NEW populations using the ratio of the residual to additive genetic variance of the REF population and assuming that the genetic variance is entirely additive, polygenic, and fully captured by the markers. Additionally, as long as the potential parents of the NEW population is known and their marker genotypes are phased, genotypes of potential NEW populations can be simulated using software like `AlphaSimR`(Faux et al., 2016; Gaynor et al., 2021).

Using classic results for predicting the accuracy of BLUPs Henderson (1975), and as derived in the Appendix, Figure 4 shows that the predicted accuracies calculated using only the phased genotypes of the REF population and an estimate of 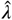 are generally less biased and more precise than the corresponding cross-validation-based estimates, particularly for Diverse REF populations. Predicted accuracies in diverse populations were most accurate when the genetic architecture was polygenic and driven by common variants, as expected because the predictions assume this architecture. But even when architectures were dominated by rare variants or included dominance variance that could be correlated with additive effects in the REF population, predicted accuracies remained much less biased than cross-validation estimates. In narrow REF populations, cross-validation estimates were less biased than in diverse REF population, except when the magnitude of dominance variance was large, but still more biased than predicted accuracies. Interestingly, even with dominance, the analytical predictions remained fairly accurate.

**FIGURE 4.**
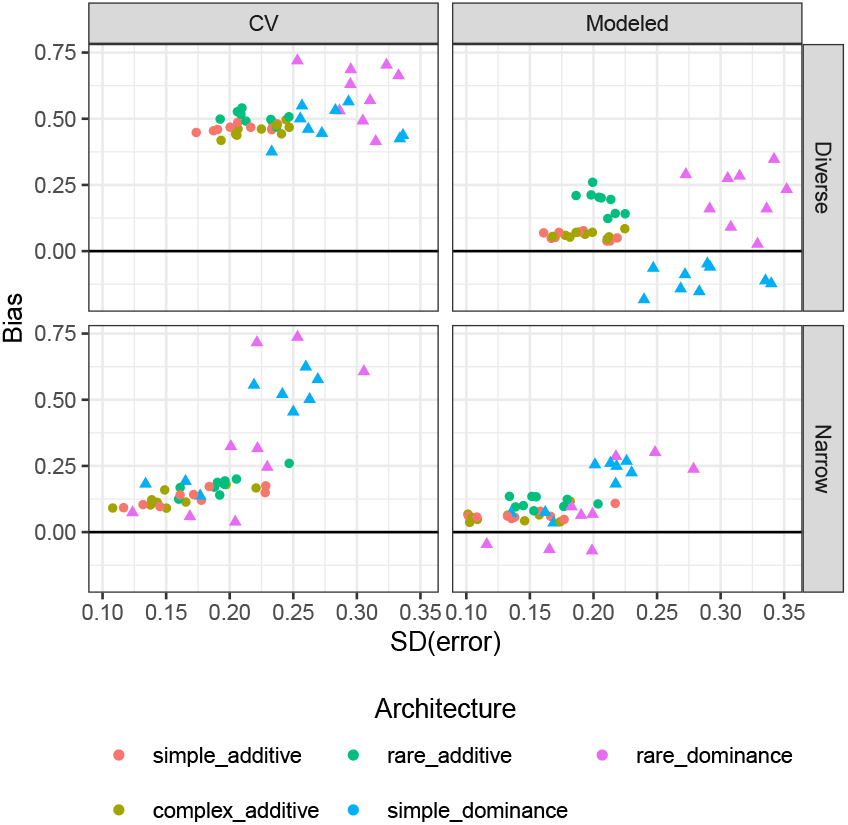
Analytical predictions are more reliable than cross-validation for evaluating REF populations for Recurrent Genomic Selection. Using the same simulations as above, analytical predictions of accuracy were made using only the estimate of 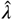 from the phenotype data of each REF population and genotypes from the simulated cross of two selected parents (10% selected based on phenotype). Cross-validation estimates of accuracy and analytical predictions were compared to the actual accuracy in these simulated individuals. Bias (mean error), and the standard deviation of errors were calculated across simulations with the same REF population, architecture, and estimate of predictive ability from cross validation.

### Geometry of Genomic Prediction accuracy in future generations

Beyond providing more accurate estimates of GP accuracy than cross-validation, the mathematical formula of GP accuracy used in Figure 4 is illustrative of *why* GP accuracy differs between the current and future generations for some REF populations more than others. As derived in the supplements, under an additive genetic architecture with known QTL, and when a GBLUP or snpBLUP model is used for Genomic Prediction, GP accuracy is a function of i) the Principal Components (PCs), or Singular Value Decomposition of the genotype matrix of the REF population, ii) the ratio of environmental noise to genetic variance, and iii) the genotype matrix of the future (or NEW) population. The steps to predict GP accuracy for any NEW population are:

1. Perform a Singular Value Decomposition of the REF genotype matrix and extract each of the right singular vectors, (*i.e*., the eigenvectors of the linkage disequilibrium (LD) matrix) of the REF population. These are the genomic features that GBLUP uses to make predictions.
2. Calculate the accuracy of the GP model along each feature, which is a function of the variance of the REF population along the corresponding eigenvector and the signal-to-noise ratio *λ*. Call this *w*_*i*_.
3. Simulate the genotypes of a relevant NEW population by simulated crosses among selected parents.
4. Measure the variance of the NEW genotypes when projected along each of these eigenvectors. Call this 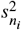. Let 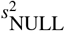 be the variance of the NEW genotypes that doesn’t lie along any of the eigenvector features.
5. Using the set of *w*_*i*_ and 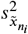 for each of the genomic features, predict the GP accuracy as:

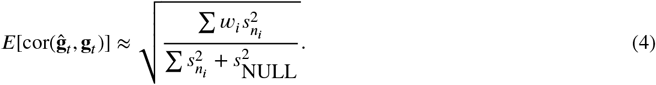

Thus, GP accuracy can be understood by studying the three parameters: 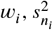, and 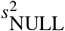.

Starting in reverse, let 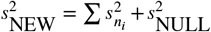 be the total genotype variance in the NEW population, which is proportional to the total genetic variance under our assumption that QTL effect sizes follow a Gaussian distribution and all genetic variance is additive. Thus, 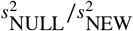 is the fraction of genotype variance that is *inaccessible* to a GBLUP model, regardless of the precision of the REF phenotype data, simply because this variation does not segregate in the REF population. For example, two loci that are in perfect LD in the REF population but become separated by recombination in the NEW population would increase 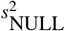. To demonstrate, I simulated 10 full-sib families made from crossing two randomly selected inbred lines from each of the six REF populations. Within crosses derived from the Diversity_212 population, ≈ 75% of variation was completely inaccessible to a GBLUP model (Figure 5A). In contrast, within crosses derived from the Biparental_212 or NAM_566 populations, only ≈ 25% of variation would be inaccessible. 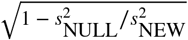 thus sets the upper bound of the GBLUP accuracy in the NEW population.

**FIGURE 5.**
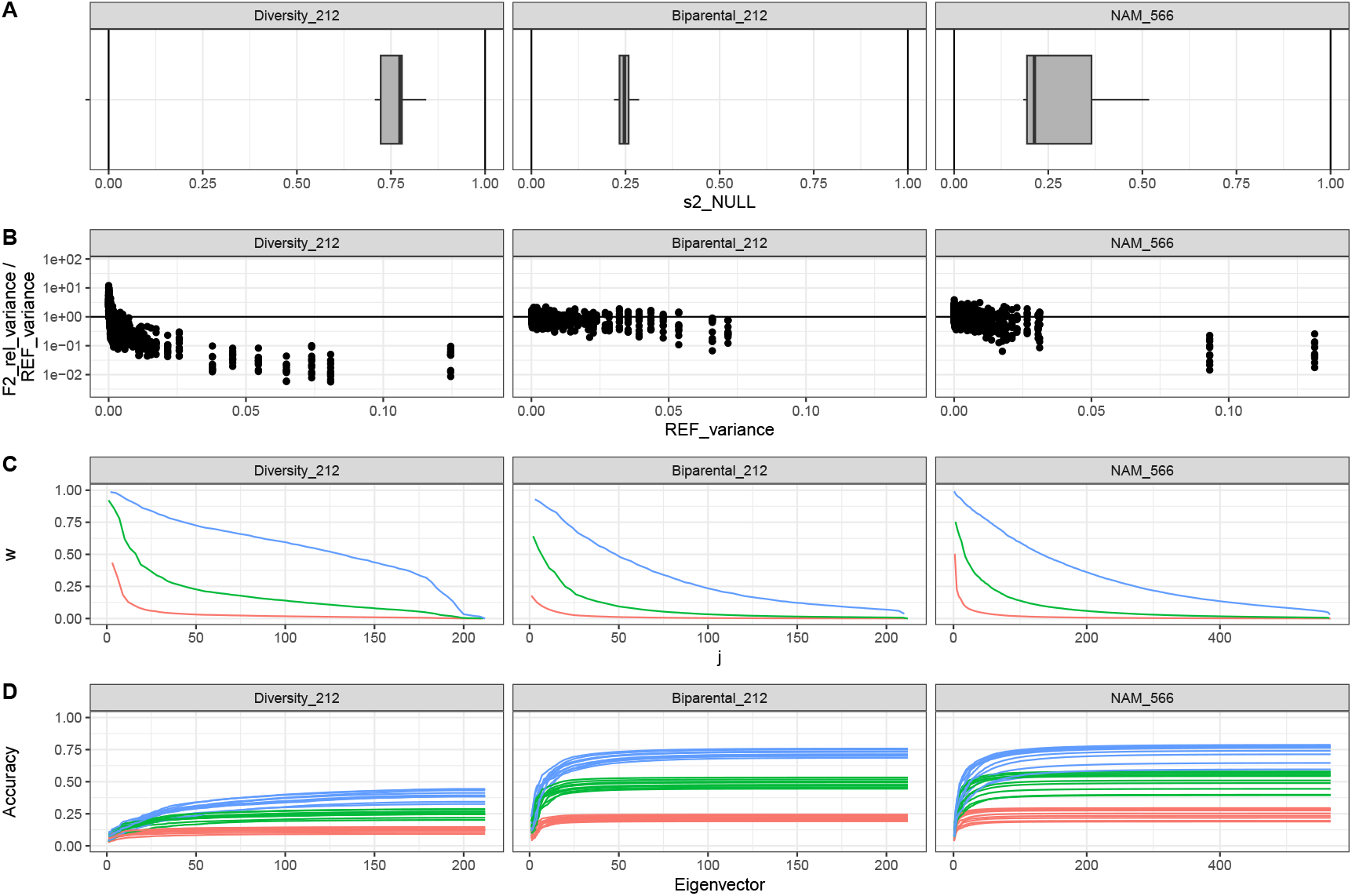
Determinants of Genomic Prediction accuracy using GBLUP for three reference populations. Three reference populations are shown: a Diverse population of size 212, a biparental F2-derived DH population of size 212, and a 4-parent NAM of size 566. For each population, decompositions of GBLUP accuracy are demonstrated for 10 target populations, each composed of the F2 progeny of the cross between two randomly sampled parents from the reference population. A) GBLUP null-space: The percentage of variance in the target population completely inaccessible to the GBLUP model. B) Eigenvalue relative-importance between target and reference populations. For each eigenvalue of the reference population, its size (i.e., variance in the reference population) is plotted against its relative importance between the target and reference populations. Points below the y=1 line denote eigenvalues with less importance in the target population than in the reference population. C) GBLUP feature accuracy. GBLUP models are predictions using the eigenvectors of the reference population as features. For each eigenvector feature, the w statistic gives its weight, or squared prediction accuracy. w values are a function of the variance in the training data (i.e., eigenvalue), and the signal:noise term *λ*, and are shown for three levels of residual variance in the training data, corresponding to *h*^2^ = 0.1 (red), 0.5 (green), or 0.9 (blue) in the diverse population. D) Accuracy of GBLUP models using the first *k* eigenvector features (x-axis), given the same three levels of residual variance in the training data.

Next, 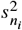 is the variance within the NEW population along the *i*th PC of the REF population. PCs are sorted in order of decreasing importance in the REF population. High-ranked PCs do not necessarily capture the most variance in the NEW population. Figure 5B shows the ratio of the proportion of variance explained by each PC in the NEW population relative to the REF population. Within crosses derived from the Diversity_212 population, the largest PCs capture a much lower proportion of variation than in the REF population, with only the lowest-ranked PCs capturing more variation in the NEW population. In contrast, except for the first two PCs of the NAM_566 population, most PCs capture similar proportions of variance within crosses as in the corresponding unstructured, narrow REF populations.

The proportion of NEW population variance captured by the first PCs is important because the GBLUP model can learn the genotype-phenotype associations of these PC more precisely than lower-ranked PCs, which is reflected in the *w*_*i*_ “weight” parameter. *w*_*i*_ is a function of the importance of that PC in the REF population (*i.e*., its eigenvalue) and the signal-to-noise term: 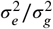 (See Appendix). Figure 5C shows weights for each eigenvector at three levels of environmental noise. Except when heritability is very high, only the first several eigenvectors have weights 0.5, meaning that the contribution of higher-order PCs to trait variation is uncertain and so will be down-weighted in genetic value predictions. Comparing Figures 5C and B, under low and moderate heritabilities, eigenvectors that capture a lot of variation in the NEW population have very low weights. Weights start lower and decline faster in the narrow populations than the Diversity_212 population because the genetic variance in the REF population is lower (meaning the signal-to-noise term is larger) and because the first eigenvectors are proportionally less important in the REF population (meaning more variance is distributed among higher-order eigenvectors).

Putting 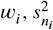, and 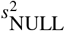 together, Figure 5D shows how GBLUP accuracy accumulates within new crosses across the eigenvector genomic features. In crosses derived from the Diversity_212 population, many eigenvector features contribute to accuracy (with meaningful contributions up to the the ≈ 175th eigenvector when *h*^2^ = 0.9, up to the ≈ 50th when *h*^2^ = 0.5, and up to the ≈ 25th when *h*^2^ = 0.1. But accuracy peaks at a low value, partially because 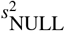 is so large, but also because the PCs that are important in the NEW population (large 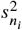) were not important in the REF population and so have low weights. In contrast, in the unstructured, narrow populations, the vast majority of accuracy is gained by the first few PCs—because they capture a large proportion of variance in the NEW population and have higher weights. Together, this gives GBLUP models trained on unstructured, narrow population much higher accuracy when applied to new full-sib families.

### Eigenvectors of the marker covariance matrix explain the discrepancy between true accuracy and cross-validation estimates

As described above, the key step in the prediction of GP accuracy for a NEW population is the projection of the NEW genotype matrix onto the PCs of the REF population’s genotype matrix. The variance of this projection for the first several PCs largely determines GP accuracy. The projection operator for the *i*th PC is the corresponding right-eigenvector of the REF population’s genotype matrix, which is a vector of weights on each marker. Figure 6 shows the marker weights for the first 10 eigenvectors of four REF populations. Except for the first two PCs in the NAM_566 population, most eigenvectors of the three unstructured, narrow populations have high weights on markers on specific chromosomes. In contrast, the first two PCs in the NAM_566 population and all PCs of the Diversity_212 population have similar distributions of weights across all chromosomes. When the NEW population is derived from crosses among individuals in the REF population, the concentration of marker weights within a chromosome is the key factor that determines the transmission of PC importance from the REF population to a NEW population. Crosses involve meioses, during which chromosomes assort independently. Therefore, genomic features with weights distributed across chromosomes contribute little to the subsequent generation, while genomic features with weights concentrated within chromosomes are “inherited” relatively in-tact. This intuition is formalized in the Supplemental Materials, where I show that the variance along a PC in a full-sib family can be modeled as the quadratic form of the marker weights for the subset of markers that segregate between the two parents of a cross and a covariance matrix determined by the probability of recombination between pairs of markers. This covariance matrix is a simple function of the genetic map; it will be close to one for marker pairs within ≈ 10cM on the same chromosome and will be zero for pairs of markers on different chromosomes. Eigenvectors with high-weight markers on multiple chromosomes will have lower quadratic form values than eigenvectors with weights concentrated on a single chromosome because a higher proportion of cross-product terms between pairs of markers will be zero.

**FIGURE 6.**
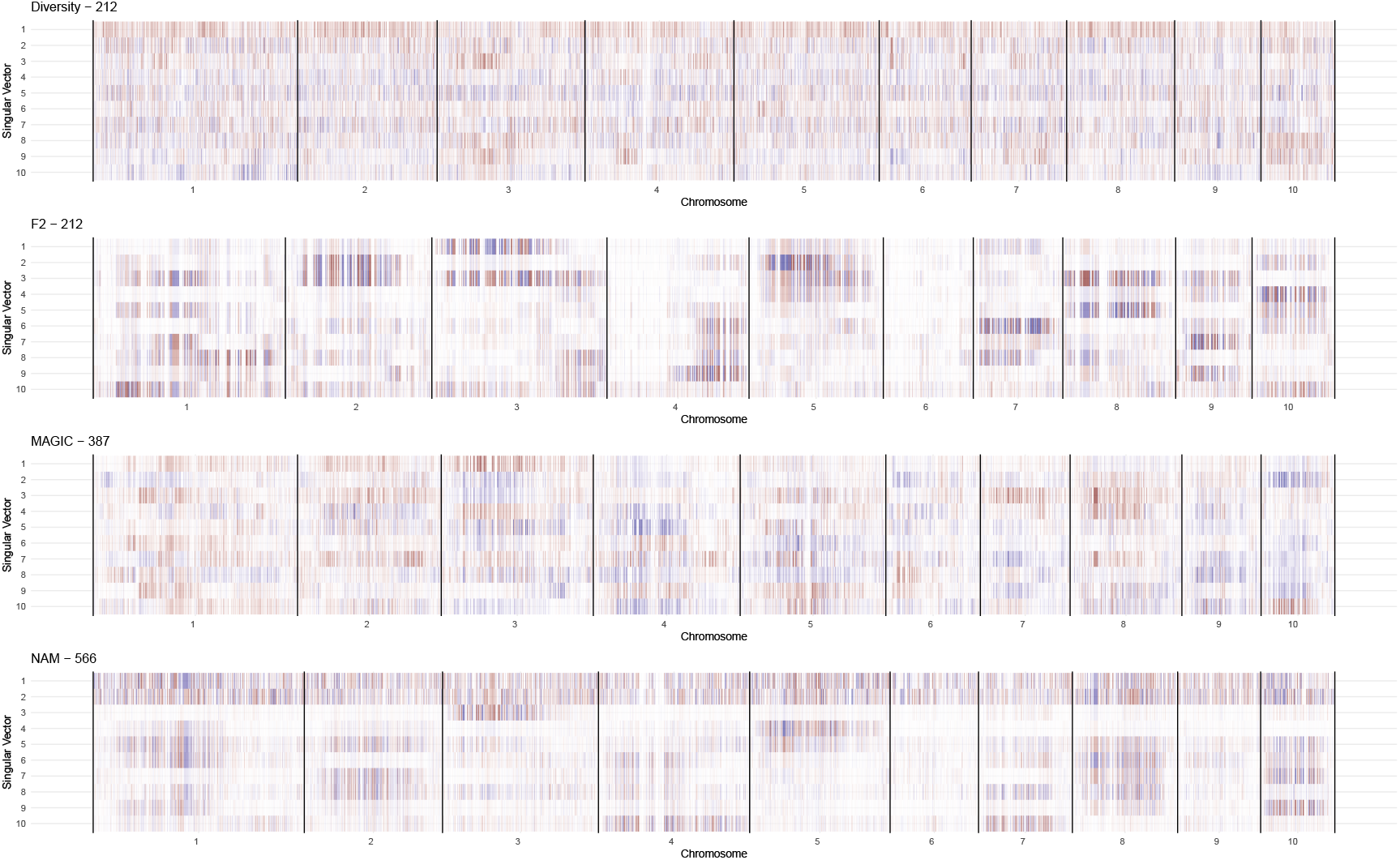
GBLUP feature projection matrices for three reference populations. GBLUP feature projection matrices are the right-singular vectors of the genotype matrix of the reference population. Multiplying these matrices by the centered genotype matrices of the target population give the scores for each feature in each target individual. The variance in these scores determines the importance of that feature in the target population. Feature projection matrices have columns corresponding to each marker, and are shown organized by position within each chromosome. Heatmaps shows weights of each marker for each projection vector, with positive (negative) weights in red (blue) for genotypes relative to the B73 reference genome. Weights close to zero are white. Only the first 10 right-singular vectors of the reference population genotype matrix are shown, as these are the only features with high weights for *h*^2^ <= 0.5 (Figure 5).

Within the REF population itself, no recombination occurs between the training and testing partition of a cross-validation scheme, so the quadratic form value will be close to 1 regardless of the distribution of marker weights across chromosomes. Thus the relative variance explained by each PC in the testing partition will be similar to that of the training partition. If the largest PCs have eigenvectors with marker weights concentrated on single chromosomes (or chromosome arms), these PCs will also be important in the NEW population and cross-validation estimates of accuracy will be reliable. But, if the first PCs have eigenvectors with high-weight markers distributed across chromosomes, these PCs will have low variance in the NEW population and cross-validation estimates will be unreliable.

Why are the marker weights along the eigenvectors of the LD matrix concentrated within chromosomes for some REF populations and distributed among chromosomes in others? The eigenvectors of the genetic relationship matrix and the eigenvectors of the LD matrix are linked through the singular value decomposition of the REF genotype matrix, as these are the left- and right-singular vectors, respectively. Conceptually, the singular vectors can be thought of as capturing causal factors for the distribution of genetic variation in a population. These causal factors could be factors causing non-random relationships among individuals (like population structure), or factors causing non-random relationships among markers (like genetic linkage and Mendelian assortment of chromosomes). For example, if individuals come from two populations, there will likely be a left-eigenvector grouping individuals by population, and a corresponding right-eigenvector representing the difference in allele frequencies among populations, a function of the divergence times and sizes of each population. Because drift is largely independent among loci, markers with large weights on this eigenvector will be distributed randomly across the genome. Because population structure affects so many markers, it tends to drive the largest PCs. Therefore, population structure in a REF population tends to result in several large PCs with corresponding eigenvectors with high marker weights distributed across all chromosomes. We see this pattern in the Diversity_212 population. Additionally, in the NAM_566 population, population structure is explained by two PCs.These two PCs have right-eigenvectors with high weights distributed across all chromosomes. However, when a REF population has little population structure, or once this structure is accounted for by higher-order PCs, the dominant remaining causal factor structuring LD will be genetic distance. Novembre and Stephens (2008) observed that when populations are distributed along a linear gradient, the eigenvectors of the genetic relationship matrix follow the basis functions of the discrete cosine transform. Markers along a chromosome are also distributed along a linear (in Morgans) gradient in recombination probability, so when recombination is the dominant causal factor driving genetic distance, the eigenvectors of the LD matrix follow the same set of basis functions, with orders and eigenvalues determined by the density of markers and the genetic lengths of each chromosome. The remaining PCs of the NAM_566 population have this characteristic. These PCs are then further structured by which markers segregate within each of the three linked biparental populations.

Therefore, in REF populations with considerable population structure, cross-validation will tend to overestimate GP accuracy for new populations proportionally to the fraction of genetic variation in the REF population captured by population-structure-driven PCs.

### Implications for research on Genomic Selection in plant breeding

Whether Genomic Prediction models will be accurate in future generations of a breeding program is perhaps the key question that will determine whether Genomic Selection can be transformative for plant breeders. If Genomic Prediction models can maintain high accuracy, then multiple cycles of selection can be made without collecting new phenotype data. In a phenotypic selection breeding scheme, phenotypic observations of each selection candidate are needed. High quality phenotype data usually requires growing candidates in multiple trials over multiple years. In many species, candidates must first be inbred for multiple generations, or made into doubled haploid lines so that candidates can be replicated in sufficient numbers to grow plots at an agronomically relevant planting density. Each of these steps takes significant time, meaning that each cycle takes multiple years. In contrast, rapid cycling strategies for many species (including wheat, maize, and rice) are available that can grow plants and make crosses multiple times per year. Since relevant phenotype data is impossible for these rapid cycling plants, selections can only be made by Genomic Prediction, and so will only be successful if the Genomic Prediction model can accurately distinguish among the candidates of the current generation.

My simulation results suggest that Genomic Prediction models trained on diverse populations with moderate population structure are unlikely to be very accurate in future generations, limiting the potential for Recurrent Genomic Selection schemes to realize its full potential. These results contrast strongly with the impression of Genomic Prediction accuracy provided by a sampling of the plant breeding literature over the past 20 years, which typically report relatively high empirical estimates of Genomic Prediction accuracy. I believe that these empirical estimates are providing a misleading picture of the potential for Recurrent Genomic Selection in plant breeding. Cross-validation-based estimates of GP accuracy do not separate accuracy into the components that persist across generations and the components that will not. However, I showed an approach to analytically decompose these components of accuracy for GBLUP / snpBLUP Genomic Prediction models. The persistence of within-family GBLUP accuracy across generations is easily understood based on the relative importance of genetic features that are chromosome (or sub-chromosome)-specific. Pan-genomic features (typically caused by population structure) contribute little to the genetic variance within families in future generations. Thus, models that gain their accuracy from such features will lose accuracy in the future. In contrast, sub-chromosomal features are largely inherited intact (if they segregate in a specific cross), and thus models that can precisely estimate the effect sizes of such features will maintain accuracy.

This intuition provides a straightforward test of the suitability of a specific breeding population for implementing Recurrent Genomic Selection, even before collecting the high-quality phenotype data needed to start making selections—the fraction of genetic variance captured by pan-genomic features sets an upper-bound on the potential accuracy of Genomic Prediction models in future generations. This intuition may be helpful for designing breeding populations that are optimal for Recurrent Genomic Selection, a topic I will explore in a follow-up paper.

### Comparison with other expressions for Genomic Prediction accuracy

A number of authors have studied Genomic Prediction accuracy analytically (Daetwyler et al., 2008; Goddard, 2009; Hayes et al., 2009; Rabier et al., 2016; Dekkers et al., 2021). Key parameters of these models include the REF population size, the trait heritability, the genetic architecture (i.e., number and effect sizes of QTL), LD between markers and QTL, and the relationship between the REF and NEW population. As I do here, most models assume that traits of breeding interest are highly polygenic, so the number of QTL is vary large and distribution across the genome. However, QTL tend to be in LD in any population, due to selection or pedigree structure. Rather than counting the number of QTL explicitly, most models use a parameter called *M*_*e*_ to represent the “effective number of independently segregating QTL”, which is more closely related to GP accuracy than the actual number of QTL. While a number of strategies for predicting or estimating *M*_*e*_ have been proposed, predictions of GP accuracy tend to be highly sensitive to *M*_*e*_ (Brard and Ricard, 2015). Also, with the exception of Wientjes et al. (2016) and Dekkers et al. (2021), these equations mostly apply to only REF populations of unrelated individuals, or only single full-sib or half-sib families (Hayes et al., 2009).

My approach for predicting GP accuracy more closely resembles that of Rabier et al. (2016) and Rabier and Grusea (2021), which draw on classic properties of linear mixed models (VanRaden, 2008). Instead of trying to estimate *M*_*e*_ and assuming unrelated individuals, I focus on the eigenvalues of the estimated Genomic Relationship matrix **K**_*r*_. The magnitude and rate of decay of eigenvalues can be compared to 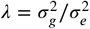 to predict accuracy in the REF population, similarly to how *M*_*e*_ and *h*^2^ interact in the earlier analytical approaches. The projection of the NEW population’s genotypes onto the eigenvectors of the REF population’s genotypes is related to the number of “effective” QTL that would segregate in the NEW population. However, it is important to stress that eigenvectors should not be interpreted mechanistically as QTL. Eigenvectors are simply structures constructed to efficiently represent the variation in the REF population and do not cleanly represent either genetic mechanisms or genetic inheritance. Nevertheless, as I show in Figure 6, the eigenvectors of the LD matrix in the REF population can be inspected for intuition as to whether or not the genetic features that explain the most variation in the REF population are likely to be important in a NEW population derived from crosses among REF population parents.

This is the place where the approach that I show here seems to be the most novel. While the above studies have provided great insight into general properties of Genomic Prediction, they have remained largely agnostic to actual relationships between REF and NEW populations. In plant breeding programs, NEW populations will generally be full-sib families derived from biparental crosses among existing lines. As long as phased genotypes of the parental lines are available, it is easy to simulate highly realistic full-sib offspring genotypes. Thus, the approximations of LD decay used by Dekkers et al. (2021) are not necessary; simulation-based estimates of this decay are simple to generate, and when combined with analytical predictions of GP accuracy for the simulated genotypes, breeding program optimization can be done very efficiently.

### Alternative strategies for measuring the utility of a Genomic Prediction model

The reason that cross-validation remains so popular as a way to assess Genomic Prediction models is that it purports to not rely on assumptions about genetic architecture (Daetwyler et al., 2013; Legarra and Reverter, 2017). Unfortunately, as I demonstrate here, cross-validation is typically used in a way that produces an unbiased estimate of the wrong quantity, relative to breeding system design. Cross-validation using random REF/NEW partitions of a population produces an estimate of the accuracy of a Genomic Prediction model for the *total genetic values* of additional candidates of the same population that have not yet been phenotyped. Accurately predicting total genetic values of these individuals will usually not provide much benefit to a breeding program. Selecting among a larger population can increase selection intensity, but gain is approximately logarithmic with intensity, so very large increases in population size would be needed for meaningful benefits, and genotyping costs remain too high to genotype very large candidate populations. Additionally, if there is significant non-additive genetic variation, the correlation of predictions with actual phenotypes of candidates themselves can be biased by this non-additive variation, and may not translate to gains in future generations. As pointed out by Dekkers et al. (2021) and Legarra and Reverter (2017), cross-validation is not reliable as an estimate for the accuracy of the genetic values of the candidates of the REF population themselves (because they do have available phenotype data), and as I showed in Figure 3, cross-validation is not reliable as an estimate of the accuracy for future populations where significant genetic gains could be made.

I propose instead to use analytical expressions for the expected Genomic Prediction accuracy. The problem with relying on analytical expressions for Genomic Prediction accuracy is that if the true genetic architecture differs too much from the idealization represented by a GBLUP model, the expressions may be unreliable. While analytical accuracy estimates did decline for non-additive architectures in my simulations (Figure 3), they remained much more reliable than cross-validation estimates. Nevertheless, if neither analytical estimates nor cross-validation is acceptable, what strategies could practitioners use to critique Genomic Prediction models?

Ultimately, I believe that Genomic Prediction models should be evaluated based on their ability to accurately predict the ranking of full-sib candidates based on the average performance of the offspring of these candidates when crossed randomly to other candidates. Progeny testing like this is the way that genomic evaluations in animal breeding are evaluated (Legarra and Reverter, 2018) and are the only way to directly measure additive genetic values. Doing progeny testing for future candidates would be a tremendous investment for plant breeding programs, requiring at least two new generations (plus potentially multiple rounds of selfing for line development) to be created and intensively phenotyped to get accurate estimated breeding value estimates. However, converting a breeding program to Genomic Selection is also a major investment, and shouldn’t be undertaken if the benefits are not clear.

Alternatively, the goal of using Genomic Prediction models in breeding is to increase the rate of genetic gain. Since its invention more than 20 years ago, there have been several historical studies demonstrating increased rates of genetic gain in cattle using Genomic Selection (García-Ruiz et al., 2016; Doublet et al., 2019; Wiggans and Carrillo, 2022). Measuring genetic gains requires “era” studies that directly compare progenitor and new varieties, ideally produced from programs using different selection strategies. I have found only a few examples of plant breeding studies that have measured gains from Genomic Selection. Some examples include: (Beyene et al., 2015; Vivek et al., 2017; Zhang et al., 2017; Herter et al., 2019; Das et al., 2020; Bonnett et al., 2022). The reported realized genetic gains in these studies are mixed, with some showing improvements over conventional methods and others not. But in all cases, these are very short-term studies, so the long-term benefit of Genomic Selection in each case is unclear. Therefore, more long-term studies are needed to demonstrate the usefulness of Genomic Selection and Genomic Prediction models specifically in plant breeding.

## Conclusions

With this collection of simulations, analytical results, and literature surveys, I aim to establish four main points:

1. The same Genomic Prediction model trained on the same population with the same phenotype data will have a different accuracy for different target populations. It is trivial to increase accuracy by increasing training population size, phenotyping intensity, or altering the target population composition, if these are not constrained by practical limitations of real breeding programs, and if the objectives are only accuracy, not genetic gain. Thus, accuracy values reported in isolation, without clearly specified parameters tied to real breeding program constraints, and without clearly links to genetic gain, have little value for evaluating Genomic Selection as a plant breeding tool.
2. Recurrent Genomic Selection (without model retraining) could be the most impactful use of Genomic Selection in breeding. Within-family Genomic Prediction accuracy is the key metric for determining if Recurrent Genomic Selection will be successful. Accuracy measured on populations with structure is not informative for evaluating Recurrent Genomic Selection.
3. The vast majority of the plant breeding literature has not evaluated Genomic Prediction models in a way that is informative about the potential success of Recurrent Genomic Selection. Instead, the use of cross-validation, especially cross-validation evaluated on testing populations with structure, tends to produce estimates of Genomic Prediction accuracy that are greatly inflated relative to the accuracy achievable in a relevant Recurrent Genomic Selection scheme. This gives the impression that Recurrent Genomic Selection will work in many cases where it will not.
4. Phenotype data is helpful, but is not necessary to determine if Recurrent Genomic Selection can be successful for a potential breeding population. The eigenvalues and corresponding right-eigenvectors of the population’s genotype matrix are sufficient to reliably predict within-family GP accuracy and thus evaluate the potential for success using Recurrent Genomic Selection.

Finally, I suggest that the plant breeding community should stop relying on cross-validation for evaluating Genomic Prediction, except under limited narrow contexts, perhaps including CV2 and CV0 breeding scenarios where the goal is closer to phenotype prediction than breeding value prediction. Instead, analytical calculations of accuracy, or progeny testing, are more appropriate tools for evaluating Genomic Prediction models for use in Genomic Selection breeding schemes.

## Author Contributions

## Supplemental Materials

## Supplemental Figures

**SUPPLEMENTAL FIGURE S1.**
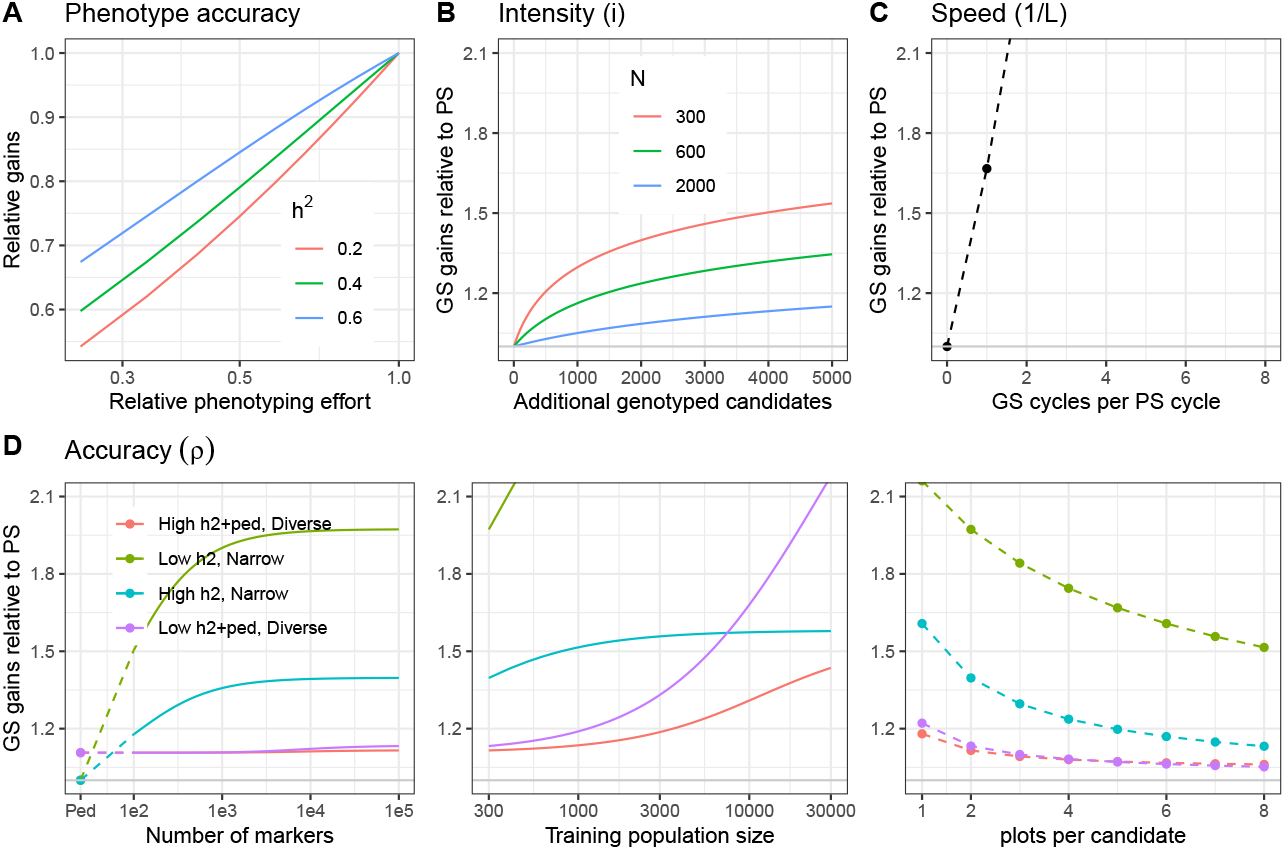
An analysis of the breeder’s equation illustrates why the most impactful use of Genomic Prediction is to enable faster cycles. **A**. Impact of reducing phenotyping intensity on rates of genetic gain using phenotpic selection for three levels of full-intensity *h*^2^. I assume reducing intensity occurs from running fewer trials per candidate, with accuracy using trials being 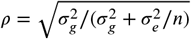. **B** - **D** are the same as Figure 1, but with expanded y-axes. **B** Rates of genetic gain when using GP to evaluate additional candidates of the Cycle 0 generation without phenotype data, relative to phenotypic selection for three sizes of populations of phenotyped candidates as a function of the number of additional candidates evaluated by GP. **C** Rates of genetic gain when using GP to evaluate candidates of future cycles, eliminating the need to wait for phenotype data, *i.e. Recurrent Genomic Selection*, relative to phenotypic selection. Values assume PS cycles require 5 years and GS cycles can occur 4x per year. **D** Rates of genetic gain when using GP to improve the accuracy of the reference population of candidates in Cycle 0, *i.e*., those with phenotype data, relative to phenotypic selections of these candidates. Three ways to increase the intensity of GP are evaluated: i) increasing the number of markers genotyped, ii) increasing the reference population size, and iii) improving the quality of the phenotype data by higher replication per candidate. In each panel, four types of reference populations are used, varying in genetic diversity (Diverse vs Narrow), heritability (*h*^2^), and whether a pedigree is available and informative (+ped). Exact parameters are given in the Methods. A zoomed-in version of the figure is available as Supplemental Figure **??**.

### Simulation of Recurrent Genomic Selection

I designed a simple simulation of a Recurrent Genomic Selection (RGS) breeding scheme to demonstrate that within-family accuracy is required for a Genomic Prediction model to be useful; in particular that accuracy in future populations with structure does not guarantee successful outcomes.

I simulated a population of 20 outbred diploid candidates segregating for 10 QTL with allele frequencies ≈ 0.5, a population where GP could have very high accuracy. Each QTL was placed on a separate chromosome, and thus all were unlinked. Then I created 10 SNP markers, each in perfect LD with one of the QTL. These SNPs were either placed adjacent to their paired QTL on the same chromosome, in which case markers and QTL stay in perfect Linkage Disequilibrium (LD) across generations (assuming no mutation), or they were placed on 10 additional chromosomes, in which case markers and QTL assort independently during meiosis and thus are uninformative about genetic values within families. However, in the latter case, some LD between markers and QTL persists across the whole population for several generations of random mating. Thus, markers do remain informative for the whole population.

**SUPPLEMENTAL FIGURE S2.**
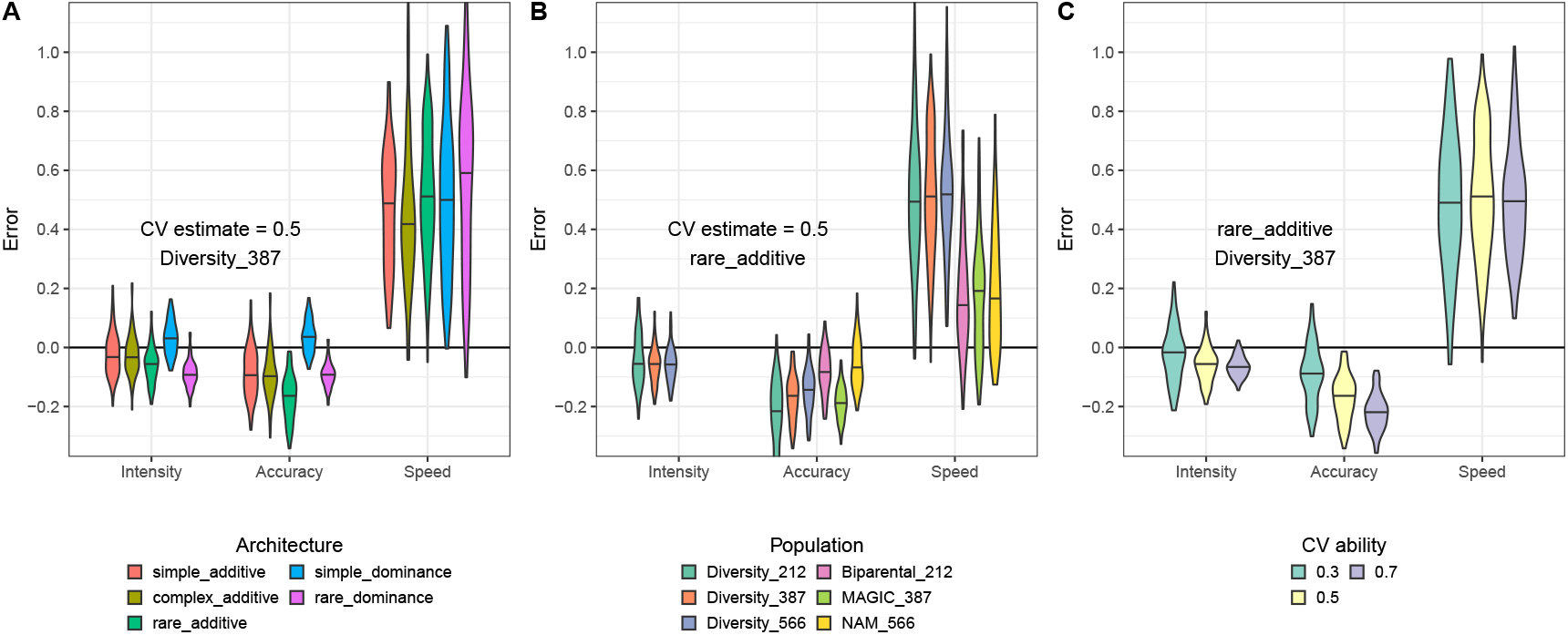
Cross-validation estimates of accuracy have higher average errors when used to evaluate Genomic Prediction accuracy. Each plot shows the density of errors 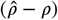 when 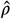 is estimated using cross-validation as 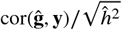 using the correction of predictive ability to accuracy of Legarra et al. (2008), for NEW populations relevant to increasing intensity, accuracy, or speed. Violin plots show densities of 100 simulations per setting. **A**. Simulations for the Diverse_387 population when cross-validation estimates predictive ability as *r* = 0.5 for five different genetic architectures. **B**. Simulations for six different REF populations when cross-validation estimates predictive ability as *r* = 0.5 using the `rare_additive`genetic architecture. Note that for the three narrow populations, no additional relevant hybrids exist, so it is not useful to use GBLUP to increase intensity. **C**. Simulations for the Diverse_387 population with a `rare_additive`genetic architecture, for different estimates of predictive ability by cross-validation.

Using these artificial individuals, I simulated 5 cycles of Recurrent Genomic Selection:

1. I simulated phenotype values for the 20 candidates assuming a perfectly additive genetic architecture with *h*^2^ = 1. I used these phenotypes to train a Genomic Prediction model using *AlphaSimR*’s *RRBLUP* function (Gaynor et al., 2021).
2. For each recurrent selection generation, I selected *n*_*s*_ ∈ *c*(1, 2, 5, 10) candidates based on their predicted genetic value. I crossed these selected candidates in all combinations, including selfing, expanding the population to a population size of 1,000.
3. In each new generation, I measured the average true genetic value of the current population, the overall selection accuracy as the Pearson Correlation Coefficient (PCC) between predicted and true genetic values across all individuals, and the average PCC within each of the individual families.

I repeated the whole simulation 1000 times for each of the two marker-QTL linkage scenarios, and reported the average values for genetic gain and within-family and across-population genomic prediction accuracy across simulations.

Figure S4 shows that when markers and QTL were adjacent, RGS resulted in genetic gains for 2-5 generations. Increasing the number of candidates selected each generation from one to 2-10 resulted in lower initial rates of gain (because candidates 2-10 had lower genetic values than the top candidate), but positive rates of gain persisted longer, resulting in slightly better long-term gains.

**SUPPLEMENTAL FIGURE S3.**
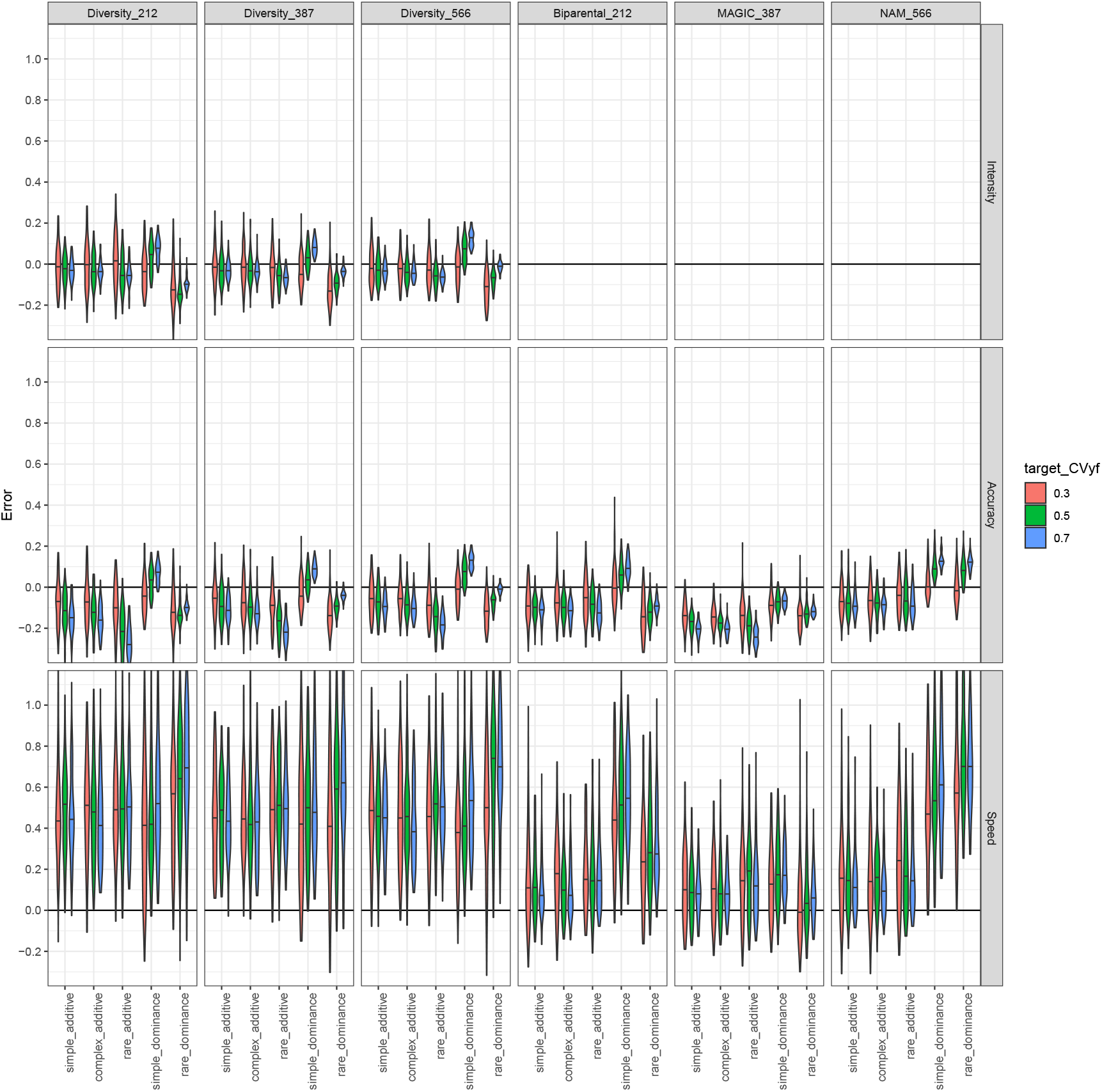
Cross-validation estimate errors differ among REF populations and genetic architectures. Each plot shows the density of errors 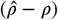 when 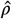 is estimated using cross-validation as 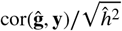 using the correction of predictive ability to accuracy of Legarra et al. (2008), for NEW populations relevant to increasing intensity, accuracy, or speed. Violin plots show densities of 100 simulations per setting.

**SUPPLEMENTAL FIGURE S4.**
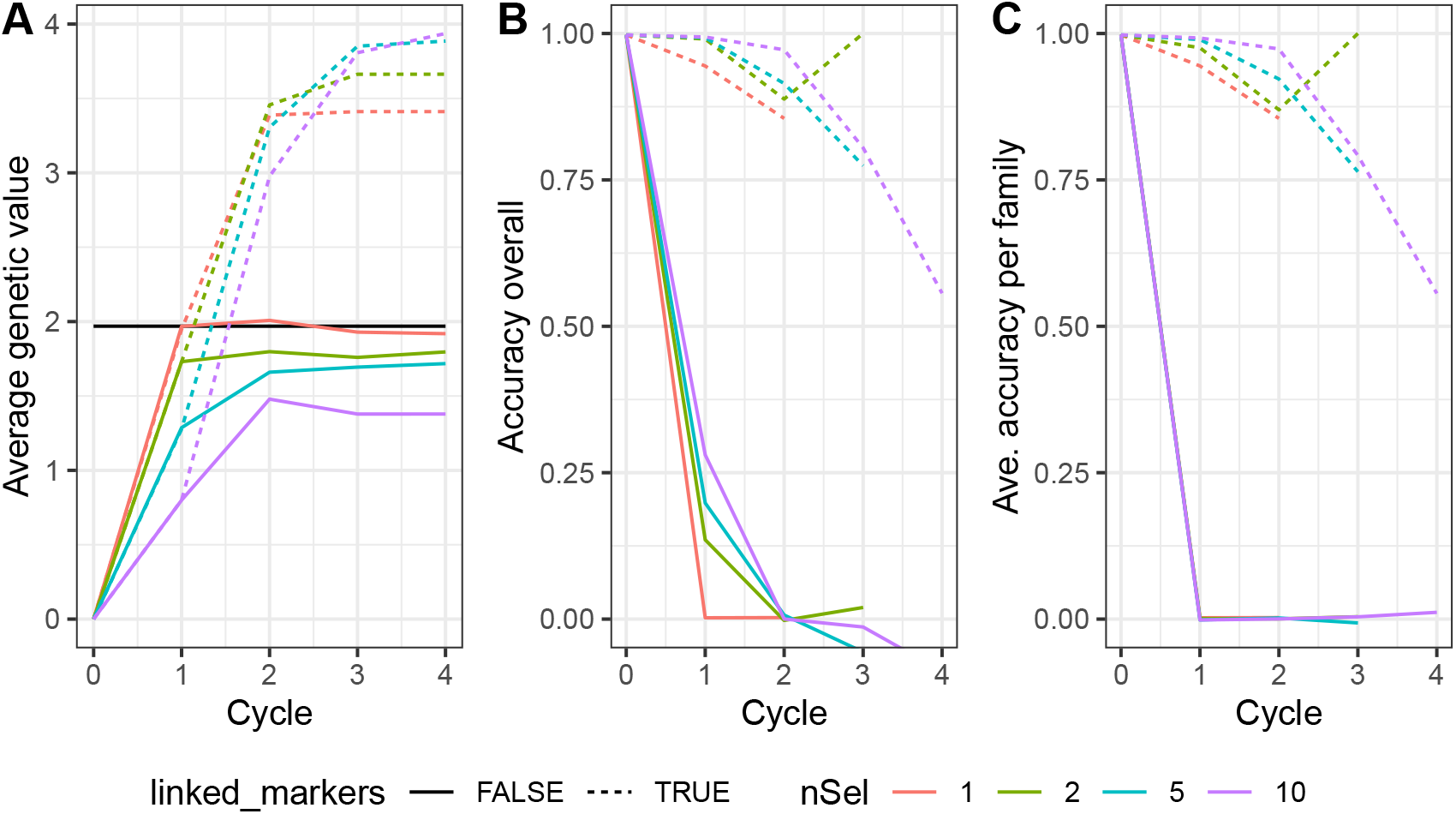
Recurrent Genomic Selection (RGS) requires within-family Genomic Prediction (GP) accuracy. Simulations of four cycles of RGS in situations where the GP model had high (*linked_markers* = TRUE, dashed lines) or zero (*linked_markers* = FALSE, solid lines) within-cross accuracy. The first cycle selected among 20 founder candidates with phenotype (and genotype) data based on estimated genetic values. In cycles 2-4, selections were based on predicted genetic values using the GP model trained in cycle 1. The best 1-10 predicted individuals were selected based on the GP model in each cycle and crossed (or selfed) in all combinations for a total population size of 1000. A) Average genetic values of the candidate population, averaged over 100 simulations. B) Average GP accuracy among all candidates each generation. C) Average within-cross accuracy, averaged across all full-sib (or selfed) families.

In contrast, when markers and QTL were on different chromosomes, genetic gains were made only in the first 1-2 cycles. In particular, when only one candidate was selected, 100% of the final average genetic value was achieved in the first cycle of selection. After that, no further gain was possible because, by construction, the Genomic Prediction model had zero accuracy within full-sib families, and all candidates in the second generation were full-sibs (actually selfs of the single selected parent). In contrast, when 10 candidates were selected to form the second generation, a total of 45 full-sib families were created with 22-23 offspring per-cross. While the Genomic Prediction model had essentially zero accuracy within each of these full-sib families, it did have some ability to distinguish among these families. Thus, additional genetic gains were possible in the second generation by selecting among these families. However, the rate of gain was much lower than in the first cycle because LD between SNPs and QTL declined each cycle. Thus, while the rate of gain in the second generation was higher with *n*_*s*_ = 10 than *n*_*s*_ = 1, the total gain relative to the base generation was lower and never reached that achieved by *n*_*s*_ = 1. In fact, regardless of *n*_*s*_, in scenarios where the Genomic Prediction model was not accurate within full-sib families, the final average genetic value after five generations of RGS was never better than the genetic value of the best individual of the base population.

These results demonstrate that when the goal of breeding by RGS is to maximize the average genetic value of a population, within-family accuracy is required. Studies that evaluate GP models in validation populations composed of multiple families may show that GP models are accurate. But, this accuracy cannot result in genetic gain over simply selecting the best individual from the REF population. These results apply more generally to any type of population structure in the candidates. Therefore, studies on RGS should evaluate the accuracy of GP models in populations without structure, such as full-sib families, or at least quantify the increases in accuracy relative to the accuracy achievable using population structure or pedigree information alone.

### Derivation of approximate analytical expressions for GBLUP accuracy

Here, I derive approximate expectations for accuracy of GBLUP and snpBLUP models which I used as expressions to predict accuracy in the simulation studies of the main text. The key difference between my derivations below and earlier analytical studies of Genomic Prediction accuracy is that my calculations are for a specific target population. Earlier results focused on idealized target populations, or populations that matched the REF population in structure, which is a reasonable starting point if the target population is not yet available. While this is necessarily the case for future candidates that have not yet been created in a Recurrent Genomic Selection breeding scheme, the important genetic characteristics can be accurately simulated as long as phased genotypes of potential parents of the target population are available together with a genetic map. I use the *R* package *AlphaSimR* (Gaynor et al., 2021) to simulate individuals in relevant target programs for plant breeding schemes. While the actual candidate individuals in such real breeding programs would differ from these simulated individuals, the important characteristics—in particular the linkage disequilibrium structure among the QTL genotypes—can be accurately predicted.

I first derive an approximation for the expected accuracy conditional on phenotypes of the REF population. I then derive an approximation for the expected accuracy that would be achieved before phenotype data are available, and thus before the model can even be trained, given only genotypes and genetic and residual variances. I then show how this latter expression can be simplified using the Singular Value Decomposition of the REF genotype matrix.

In the notation below, I assume that the genetic architecture is entirely additive, and so genetic values and breeding values are perfectly correlated.

### Definitions

Let **x** and **y** be two random vectors of length *n*. var(**x**) is a scalar, the sample variance of the elements of 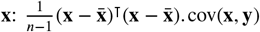 is also a scalar, the sample covariance of the (*x, y*) pairs: 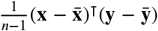. In contrast, Cov(**x, y**) is the matrix of pairwise covariance of elements of each vector across conceptual repetitive draws of pairs (**x, y**). The Pearson Correlation Coefficient between **x** and **y** is:

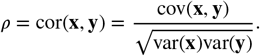

An approximation for the expectation of this correlation coefficient is:

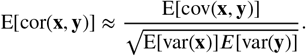

where the approximation is due to the assumption that the expectation or a ratio is approximately equal to the ratio of expectations.

This is generally accurate if *n* is large. The expectation of the covariance between **x** and **y**, the numerator of above, is:

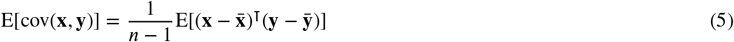

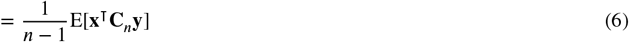

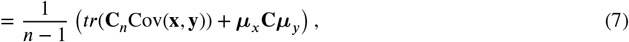

where **C**_*n*_ = **I**_*n*_ − 1/*n***11**^⊺^ is a centering matrix, tr(·) is the matrix trace, and ***µ***_*x*_ and ***µ***_*y*_ are the expected values of **x** and **y**, respectively. Thus 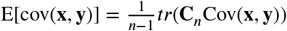 if ***µ***_*x*_ = ***µ***_*y*_ = **0**. Expectations of the variances follow similarly.

Let **X**_*r*_ be the *r* × *p* QTL genotype matrix for a reference (REF) population of size *r* used to train a Genomic Prediction model, with genotypes coded as (0, 1, 2) − 2*q*_*j*_ for *q*_*j*_ the minor allele frequency (MAF) of the *j*th locus such that columns of **X**_*r*_ sum to zero. 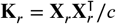 is the genomic relationship matrix (VanRaden, 2008) with 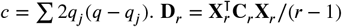 is the linkage disequilibrium (LD) matrix of the QTL genotypes for the REF population (the centering matrix **C**_*r*_ is not needed here because columns of **X**_*r*_ are centered, but is included for clarity below). Under a purely additive genetic architecture with QTL effect sizes **b**, the genetic values are **g**_*r*_ = **X**_*r*_**b**. Phenotypes of the individuals for a target trait are **y**_*r*_, with E[**y**_*r*_ ∣ **g**_*r*_] = **g**_*r*_.

Let 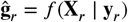 be estimated genetic values of the REF population from a Genomic Prediction model. Let **X**_*n*_ be the *n* × *p* QTL genotype matrix for a population of *n* NEW candidates, coded in the same way as **X**, and 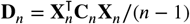 be the LD matrix. In the simulation studies of the main text, I create **X**_*n*_ by simulating crosses among specific parents with phased genotypes using *AlphaSimR* (Gaynor et al., 2021). The genetic values of the NEW population are **g**_*n*_ = **X**_*n*_**b**. Predicted genetic values are 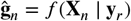.

The accuracy of the Genomic Prediction model in the NEW population is 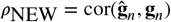. While the actual accuracy cannot be known unless **g**_*n*_ itself is known, we can study the *expectation* of its accuracy in two ways: 1) 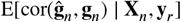, the expected accuracy given the genotypes of the NEW population and the phenotypes of the REF population, and 2) 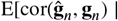 **X**_*n*_], the expected accuracy given only the genotypes of the NEW population. The first expectation will be more precise, but requires that phenotypes of the REF population are available, and thus is less useful for optimizing a breeding program. In contrast, the second expectation can be calculated with respect to different REF populations, which can help a breeder decide for which individuals to collect phenotypes. Below, I will drop the **X**_*n*_ from the expression when it is not needed for clarity.

The Singular Value Decomposition (SVD) of **X** is 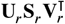, where **U**_*r*_and **V**_*r*_are orthonormal matrices and **S**_*r*_is diagonal with non-negative entries. Columns of **U**_*r*_are the eigenvectors of the genetic relationship matrix **K**_*r*_, and 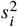, the *i*th diagonal element of 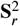 is the *i*th eigenvalue of *c***K**_*r*_. Similarly, columns of **V**_*r*_are the eigenvectors of the linkage disequilibrium (LD) matrix of the *p* QTL in the REF population (**D**_*r*_).

The GBLUP model is:

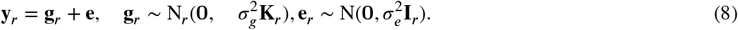

Predicted values for the REF individuals are: 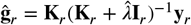 for 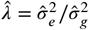. Predicted values for the NEW individuals are: 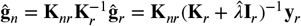, where 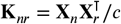. Using the SVD of **X**_*r*_we have:

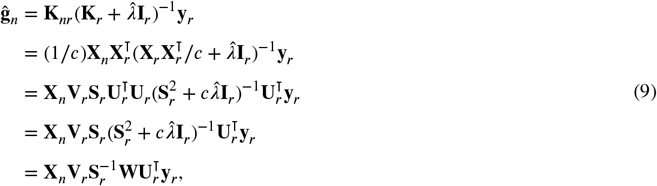

where 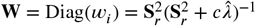, where 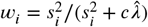 is a value between 0 and 1 for each eigenvectors of **X**_*r*_. Note that 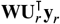 is a weighted projection of the phenotype data onto each eigenvector of **X**_*r*_, with weights given by *w*_*i*_, and 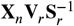 is a weighted projection of the genotypes of the NEW population onto the eigenvectors of **D**_*r*_, with weights given by 1/*s*_*i*_.

The GBLUP model is equivalent to the snpBLUP model, also known as the RRBLUP model:

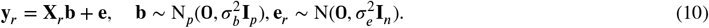

snpBLUP returns predicted values for the QTL effect sizes themselves: 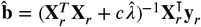, which can be used to form predicted values for the NEW individuals as 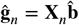. These predicted genetic values are identical to those of GBLUP. For some of the derivations below, it will be helpful to use the snpBLUP form of the predicted values.

### Accuracy conditional on phenotypes, assuming QTL and *λ* **and known**

If Equation 8 is true, such that all QTL are in **X**_*r*_, and their effect sizes are Gaussian-distributed with common variance 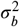, and 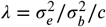 is known 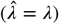, then we can compute the expected accuracy after observing phenotypes **y**_*r*_on the REF individuals. Note that this implicitly assumes that individuals in the REF and NEW populations have not undergone selection. Selection induces dependencies among QTL effect sizes and LD which invalidates the below results. Since in breeding, individuals in the REF and NEW populations have undergone selection, I evaluate the impact of this in the simulation studies in the main text.

In the case of no selection, the expectation of GBLUP accuracy given **y**_*r*_is:

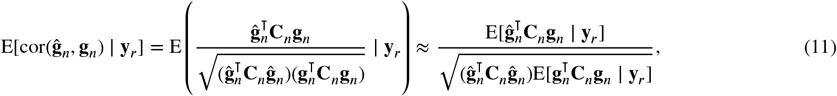

because 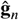 is not random given **y**_*r*_. To expand these expressions, we require the conditional distribution of **g**_*n*_∣ **y**_*r*_, assuming no selection:

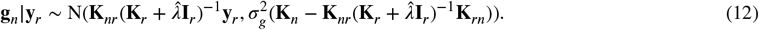

The conditional expectation and covariance of **g**_*n*_∣ **y**_*r*_can be simplified using the SVD of **X**_*r*_as:

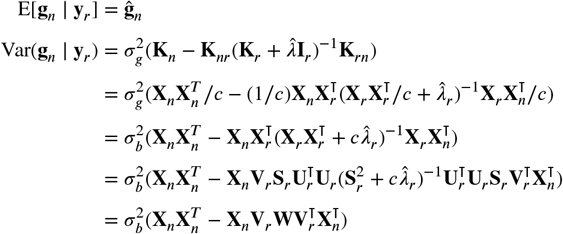

The numerator of the expected correlation can be simplified as 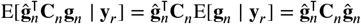. The second term in the denominator is:

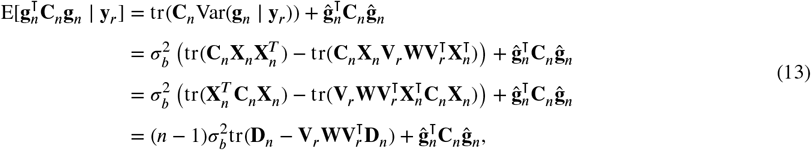

because 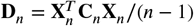. Thus, with some simplification we have:

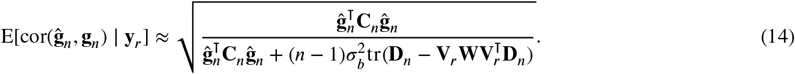

Accuracy is thus maximized when 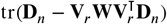 is minimized. This criterion will be discussed more below. Note that:

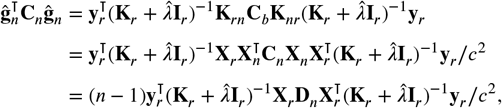

which also does not depend directly on **X**_*n*_. Thus, 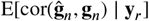 can be computed without **X**_*n*_, using only the LD matrix of the NEW individuals (**D**_*n*_) if this can be calculated without directly forming **X**_*n*_. In the case of a F2 family formed by selfing a biparental cross between two inbred lines (as used in the simulations in the main text) with genotype vectors **x**_1_and **x**_2_, and assuming the pairwise genetic distance between markers is given by the *p* × *p* matrix **R**, with distances between markers on different chromosomes set to infinity, **D**_*n*_= (1/8)(**x**_1_− **x**_2_)(**x**_1_− **x**_2_)⊺ ⊙ (**11**^⊺^ − tanh(2 **R** ), assuming the Kosambi mapping function.

### Accuracy before seeing phenotypes, assuming QTL and *λ* **and known**

Under the same assumptions as before, we can also compute the expected accuracy before seeing phenotypes of the REF population, as long as *λ* is known. In this case:

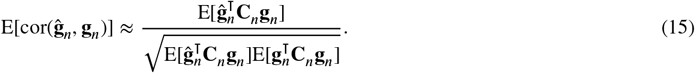

Now, both **g**_*n*_and 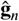, and also **y**_*r*_are random vectors with expected value equal to **0** (because I assume no selection for these derivations). The three terms of the approximation for expected accuracy can be expanded as follows:

First, we need the following expressions:

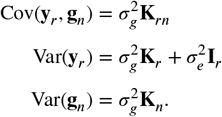

Now:

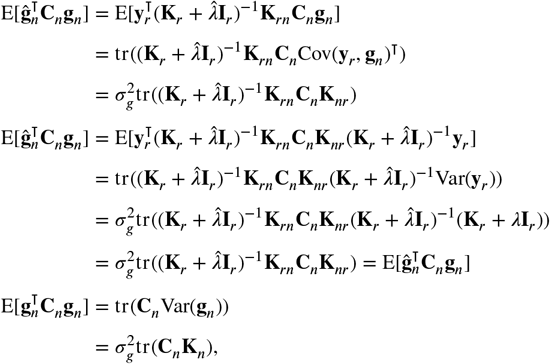

Where I assume 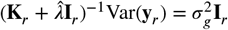 because 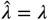. Using the SVD of **X**_*r*_, we can write these as:

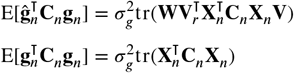

Putting this together and simplifying, we have:

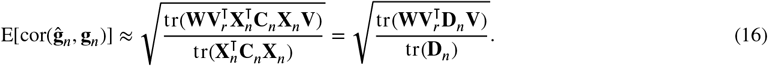

Notice that the numerator and denominator are the same as the two parts of the second term in the denominator of the approximation for 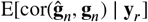 in Equation 14.

To further study this expression for unconditional expected accuracy, let 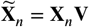 be the projection of the QTL genotypes of the NEW individuals onto the eigenvectors of the LD matrix of the REF individuals. Thus, columns of **X**_*n*_represent the coordinates of the NEW individuals along each of these axes in genotype space. Let 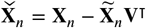, the part of the NEW genotypes that live outside the space spanned by the REF genotypes.

Using these new matrices, we can write:

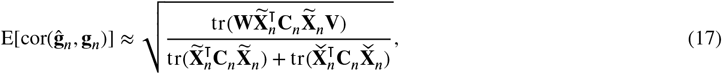

because **V**^⊺^**V** = **I**, as long as *p* < *n*, which it is in most cases. If not, **X**_*n*_= NULL and the second term will disappear.

Finally, let 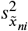 be the variance of the *i*th column of 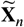, i.e., the variance of the NEW population along the *i*th eigenvector of the REF population, and recall **W** = Diag( *w*_*i*_) with ∈ [0, 1) weights for each eigenvector in the GBLUP model. The expected accuracy of a GBLUP model trained on the REF population (with these eigenvectors) in an expected NEW population, before observing phenotypes of the REF population itself, is:

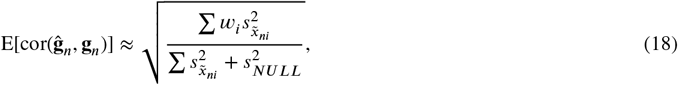

where 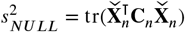 is proportional to the genetic variance in the NEW population that lies in the null space of **X** .

To interpret this result, using the singular value decomposition the variance in the QTL genotypes REF population is decomposed into a set of eigenvectors of genetic relationships (**u**_*ri*_) and LD (**v**_*ri*_), with corresponding eigenvalues 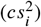. The genotype matrices of both the reference and the target population can then be represented as a sum of scores along each of these dimensions of genetic variation. In the reference population, the variance along each axis is proportional to 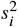, the squared singular value. It is these variances that allow the GBLUP model to learn how to predict genetic values. The model’s accuracy along each of these axes is given by 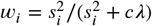, which approaches one as 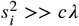, and approaches zero as 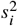 decreases for higher-order singular values. Note that the sum of 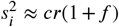, where *f* is the average inbeeding coefficient in the reference population. Thus in the expression for *w*_*i*_, the *c* terms cancel (but *r*(1 + *f* ) does not), so *w*_*i*_is approximately unaffected by the number of QTL.

In the NEW population, the variances of the candidates along each eigenvector may differ from those in the reference population. Since the first eigenvectors are defined as those that maximize the variance in the REF population, these will likely explain less variance in the new population, with other higher-order eigenvectors contributing relatively more. The more distantly related the NEW population is from the REF population, the less the eigenvectors will contribute in the new population. Higher-order eigenvectors have smaller weights (*w*_*i*_) because the 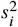 are smaller. Therefore, in most cases accuracy in the NEW population will be lower than in the REF population, with the extent of the decrease determined by how small *w*_*i*_is for the eigenvectors that capture the variance in the new population.

